# A long-term ecological research data set from the marine genetic monitoring programme ARMS-MBON 2018-2020

**DOI:** 10.1101/2024.09.26.614897

**Authors:** Nauras Daraghmeh, Katrina Exter, Justine Pagnier, Piotr Bałazy, Ibon Cancio, Giorgos Chatzigeorgiou, Eva Chatzinikolaou, Maciej Chełchowski, Nathan Alexis Mitchell Chrismas, Thierry Comtet, Thanos Dailianis, Klaas Deneudt, Oihane Diaz de Cerio, Markos Digenis, Vasilis Gerovasileiou, José González, Laura Kauppi, Jon Bent Kristoffersen, Piotr Kukliński, Rafał Lasota, Liraz Levy, Magdalena Małachowicz, Borut Mavrič, Jonas Mortelmans, Estefania Paredes, Anita Poćwierz-Kotus, Henning Reiss, Ioulia Santi, Georgia Sarafidou, Grigorios Skouradakis, Jostein Solbakken, Peter A.U. Staehr, Javier Tajadura, Jakob Thyrring, Jesus S. Troncoso, Emmanouela Vernadou, Frederique Viard, Haris Zafeiropoulos, Małgorzata Zbawicka, Christina Pavloudi, Matthias Obst

## Abstract

Molecular methods such as DNA/eDNA metabarcoding have emerged as useful tools to document biodiversity of complex communities over large spatio-temporal scales. We established an international Marine Biodiversity Observation Network (ARMS-MBON) combining standardised sampling using autonomous reef monitoring structures (ARMS) with metabarcoding for genetic monitoring of marine hard-bottom benthic communities. Here, we present the data of our first sampling campaign comprising 56 ARMS units deployed in 2018-2019 and retrieved in 2018-2020 across 15 observatories along the coasts of Europe and adjacent regions. We describe the open-access data set (image, genetic, and metadata) and explore the genetic data to show its potential for marine biodiversity monitoring and ecological research. Our analysis shows that ARMS recovered more than 60 eukaryotic phyla capturing diversity of up to ∼5,500 amplicon sequence variants and ∼1,800 operational taxonomic units, and up to ∼250 and ∼50 species per observatory using the cytochrome *c* oxidase subunit I (COI) and 18S rRNA marker genes, respectively. Further, ARMS detected threatened, vulnerable and non-indigenous species often targeted in biological monitoring. We show that while deployment duration does not drive diversity estimates, sampling effort and sequencing depth across observatories do. We recommend that ARMS should be deployed for at least three to six months during the main growth season to use resources as efficiently as possible and that post-sequencing curation is applied to enable statistical comparison of spatio-temporal entities. We suggest that ARMS should be used in biological monitoring programmes and long-term ecological research and encourage the adoption of our ARMS-MBON protocols.

## INTRODUCTION

The declining health of coastal ecosystems is a major concern for society as unsustainable use of marine resources and growing anthropogenic impacts continue to threaten marine biodiversity and ecosystem functions (Lotze et al., 2018; Smale et al., 2019; Worm et al., 2006). While the majority of coastal habitats are known to have deteriorated in the past century and to have experienced substantial declines in biodiversity (Lotze et al., 2018; Micheli et al., 2013; Obst et al., 2018), it still remains to be fully understood how the loss of species may affect the functioning of these ecosystems (Fields & Silbiger, 2022; Gamfeldt et al., 2015; Narwani et al., 2019; Virta et al., 2021). Uncertainty also prevails about the significance of individual versus cumulative anthropogenic pressures for the declining biodiversity in coastal ecosystems (Andersen et al., 2015). To better understand these relationships, consistent and well-coordinated biodiversity monitoring efforts are essential (Muller-Karger et al., 2018). These can provide comparable information on ecological variability over time and thereby help identify the thresholds at which critical changes take place (Ducklow et al., 2009). More extensive biological monitoring can also be used to contextualise taxonomic information with environmental and socio-economic data to analyse causes and impacts of biodiversity decline as well as the recovery of ecosystems in response to management and restoration efforts (Elliott, 2014; Heymans et al., 2020; Jacquemont et al., 2022) but also for following up sustainability goals set by the United Nations (UN) (e.g., Sustainable Development Goal 14: Life Below Water) and the Kunming-Montreal Global Biodiversity Framework.

To make biological monitoring programs more effective, several methods have been proposed recently (Danovaro et al., 2016), one of them being DNA metabarcoding. In principle, DNA-based techniques are capable of identifying biological communities at high temporal and spatial frequency, and with fine taxonomic resolution (Staehr et al., 2022). But despite the frequent application of metabarcoding in marine ecological research, only few marine surveillance programs have so far implemented genetic protocols for long-term monitoring (Hallam et al., 2023; Mathieu et al., 2020). Though it has been shown that DNA metabarcoding may enhance or even outperform traditional approaches (Capurso et al., 2023; Fediajevaite et al., 2021), its lack of application in large-scale spatio-temporal contexts may be due to its novelty as a biomonitoring tool (Hallam et al., 2023) or methodological biases associated with it (Capurso et al., 2023; Mathieu et al., 2020; Zhang et al., 2023). Furthermore, data management can become a demanding task when trying to link raw data with different layers of downstream analytical information across a multitude of sample localities and project collaborators, potentially discouraging the use of genomic tools in large-scale monitoring initiatives. To accelerate the use of DNA metabarcoding for long-term and geographically widespread biomonitoring, it is therefore crucial for the scientific community to develop best practices for sample collection, processing, and analysis as well as procedures and standards for data management and quality control (Santi et al., 2022).

Recently, Obst et al. (2020) established the Autonomous Reef Monitoring Structures Marine Biodiversity Observation Network (ARMS-MBON) for the genetic monitoring of hard-bottom communities, as part of the marine thematic node of the Group on Earth Observations Biodiversity Observation Network (GEO BON). This programme deploys autonomous reef monitoring structures (ARMS) in ports, marinas, and nature reserves along the European coastline, as well as at a number of locations in polar regions and the Red Sea, to capture and analyse the genetic diversity across latitudes, oceans, and benthic habitats. The use of ARMS - simple-to-produce units of stacked PVC plates mimicking the three-dimensional complexity of benthic habitats - enables standardised and non-destructive sampling of complex benthic communities (Leray & Knowlton, 2015; Ransome et al., 2017). The network maintains, to date, 25 observatories each deploying ARMS on a regular basis at one to seven sampling sites. ARMS-MBON has now become part of the European Marine Omics Biodiversity Observation Network (EMO BON) (Santi et al., 2023), a larger European initiative coordinated by the European Marine Biological Resource Centre (EMBRC) for the observation of genomic biodiversity. EMO BON includes marine biodiversity observatories from the Arctic to the Red Sea and investigates biological communities sampled from the water column and the benthic substrate using shared protocols, data, and metadata standards. Furthermore, EMO BON contributes to the UN Decade of Ocean Science for Sustainable Development by participating in the global Ocean Biomolecular Observing Network (OBON) (Meyer et al., 2022) programme, and, together with other connected projects, it aims for a worldwide coordinated biomolecular observation system (Santi et al., 2023).

Here, we present the first data release from the initial ARMS-MBON sampling campaign (i.e., from all ARMS units deployed in 2018 and 2019 and retrieved between 2018 and 2020), comprising genetic samples and image data collected by observatories from the Gulf of Finland in the Baltic Sea to the Spanish Atlantic coast, and from the northern Red Sea to the Svalbard archipelago. The data set adheres to the FAIR (Findable, Accessible, Interoperable, and Re-usable) Data Principles, and contains references to material samples, metadata descriptions, images, sequence data, derived taxonomic observations, and documentation of the analytical process. In this paper, we provide some brief exploration of the data set and give examples of potential applications, but the ARMS-MBON data can be used by any interested party for comparative studies and to support DNA-based monitoring of marine biodiversity over large spatio-temporal scales.

## MATERIALS AND METHODS

### Field sampling and laboratory protocols

The original descriptions of observatory design, field work, and sample processing procedures as well as instructions for biobanking and data management have been expanded from Obst et al. (2020). Field work and sample processing for samples of this first data release followed the official ARMS-MBON Handbook v2.0, which is in parts based on the initial protocols developed by the Global ARMS Program (https://naturalhistory.si.edu/research/global-arms-program). We used molecular protocols available under the ARMS-MBON Molecular Standard Operating Procedure v1.0 (MSOP). The Handbook and MSOP can be accessed on the ARMS-MBON GitHub repository (see Supplementary Table S1 for links). Preserved replicates of the material samples are stored and catalogued at the ARMS-MBON network partner and EMBRC member institution, the Hellenic Centre for Marine Research (HCMR), Crete, Greece, and at the individual ARMS-MBON network partner institutions.

### Access and Benefit Sharing (ABS)

All collections of ARMS samples were carried out with the necessary ABS national permits. These allow the relevant stations to collect the samples in order to utilise the genetic material for taxonomic identification purposes. If anyone wishes to obtain and process any replicate material as stored in the individual partner stations, please note that any reutilisation needs to be renegotiated with ABS competent authorities in the providing countries.

### General description of the data set for ARMS-MBON data release 001

The first ARMS-MBON data release package comprises data of 56 individual ARMS units which were deployed, retrieved and processed by ARMS-MBON network partner institutions at 15 observatories across 12 countries along the European coastline, as well as the northern Red Sea and Svalbard (Figure 1A). This covers the entirety of ARMS-MBON sampling events for which deployments took place in 2018 and/or 2019. Deployment periods ranged from 37 to 649 days during the period of April 2018 and December 2020, with the majority being deployed for around three months to approximately one year (Figure 1B). The 56 ARMS units represent 56 individual sampling events and 42 unique Unit IDs, meaning where (i) upon retrieval a new ARMS unit was deployed at the same spot for a consecutive deployment period, or (ii) for a certain period multiple units remained submerged simultaneously at the exact same spot but were either deployed or retrieved at different time points (i.e., to test the effect of deployment duration), this resulted in sampling events with the same Unit ID but separate Event IDs. General information on the observatories and ARMS deployments (e.g., coordinates, habitat type, deployment depth) and on the sampling events and their resulting material samples (e.g., date of deployment and retrieval, material sample IDs, preservative used) can be found on the GitHub repository (see Supplementary Table S1 for links) and in Supplementary Files 1 and 2, respectively.

**Figure 1.**
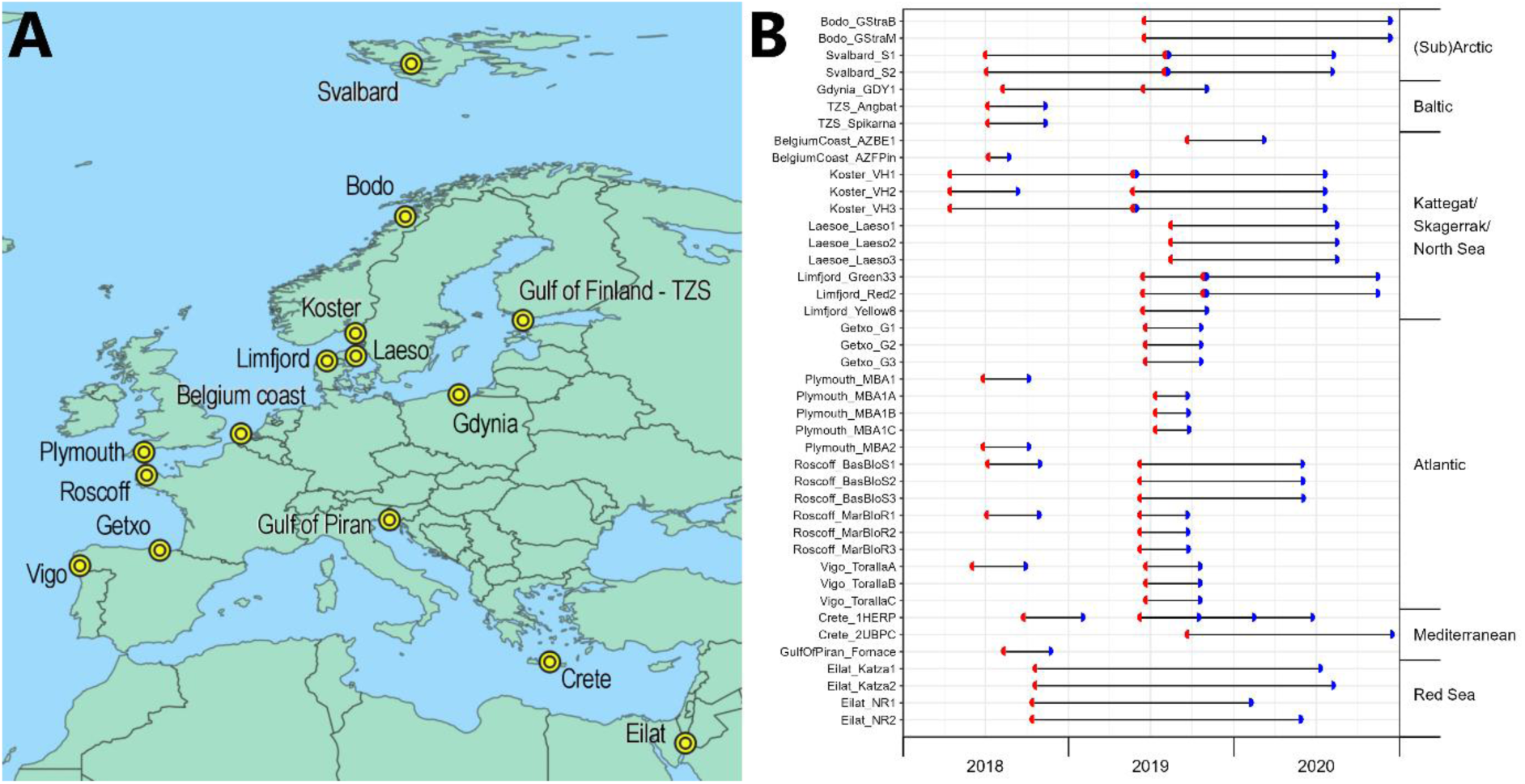
**(A)** Locations of observatories that deployed ARMS units in 2018 and 2019 during the first ARMS-MBON sampling campaign. **(B)** Sampling events of the first ARMS-MBON sampling campaign. Axis on the left shows ObservatoryID_UnitID combinations, axis on the right shows groupings of observatories into larger regions. Red semicircle: time of deployment. Blue semicircle: time of retrieval. Where red and blue semicircles meet, a new ARMS unit was deployed for a consecutive period at the same spot upon retrieval of the first unit. Where lines contain more than two semicircles, either multiple units were deployed at the exact same spot at the same time but were retrieved at different time points (see Crete_1HERP), or an additional unit was placed directly next to an already deployed one later on and both units were retrieved at the same time (see Gdynia_GDY1).

In total, this first data release package comprises data of 190 material samples. For ARMS-MBON, three size fractions (one sessile and two motile) are processed for each ARMS unit, theoretically resulting in a set of 178 biological samples for the 56 units deployed here. However, during the initial phase of ARMS-MBON, various sampling and processing techniques were tested. This resulted in (i) some biological samples being stored at −20°C without any preservative or being preserved with ethanol (EtOH) instead of dimethyl sulfoxide (DMSO; as DESS buffer solution, see Handbook), which is now the standard preservative used in ARMS-MBON; (ii) some biological samples being processed as duplicates and preserved with DMSO as well as EtOH; (iii) three sediment and two plankton samples being collected and processed as part of this sampling campaign; (iv) some sample fractions being sieved with different mesh sizes; (v) two samples being processed as technical duplicates; and (vi) only the two motile fractions being processed for one ARMS unit. Hence, a total material sample number of 190 resulted from this first campaign, including all biological and technical replicates (Table 1 and Supplementary File 1).

**Table 1.**
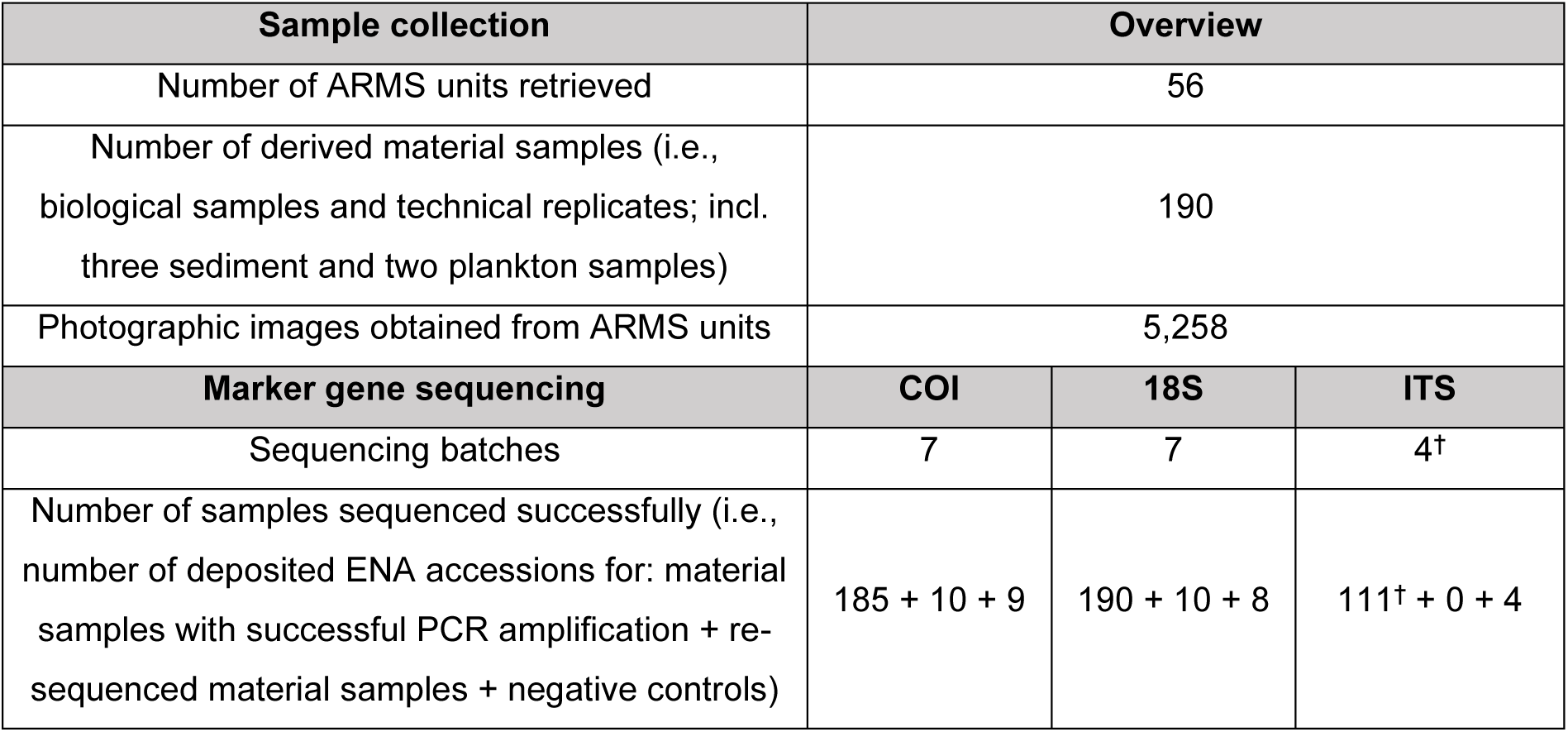
Overview of the processed samples from the first ARMS-MBON sampling campaign. † ITS amplicon sequencing was discontinued during the first sampling campaign; 111 (out of 190) samples were processed for ITS sequencing.

| Sample collection | Overview |  |  |
| --- | --- | --- | --- |
| Number of ARMS units retrieved | 56 |  |  |
| Number of derived material samples (i.e., biological samples and technical replicates; incl. three sediment and two plankton samples) | 190 |  |  |
| Photographic images obtained from ARMS units | 5,258 |  |  |
| Marker gene sequencing | COI | 18S | ITS |
| Sequencing batches | 7 | 7 | 4<sup>†</sup> |
| Number of samples sequenced successfully (i.e., number of deposited ENA accessions for: material samples with successful PCR amplification + re-sequenced material samples + negative controls) | 185 + 10 + 9 | 190 + 10 + 8 | 111<sup>†</sup> + 0 + 4 |

### ARMS plate image data

Images of each plate were recorded post-recovery and after disassembling the units, to visually document benthic communities according to the instructions in the ARMS-MBON Handbook v2.0. The image collection contains photographs of both sides of the settlement plates as well as close-ups of individual specimens or colonies. Download links for all images of this data release package are available via the GitHub repository (see Supplementary Table S1 for link and Supplementary File 1).

### Amplicon sequencing

Molecular work for samples of the first data release package was carried out as detailed in the ARMS-MBON MSOP (see Supplementary Table S1 for link) for DNA metabarcoding of the eukaryotic mitochondrial and nuclear marker genes cytochrome *c* oxidase subunit I (COI), 18S rRNA (18S) and internal transcribed spacer (ITS) region. Note that ITS was only targeted during the initial phase of ARMS-MBON and use of this marker gene has been discontinued. All samples were subjected to amplicon sequencing by the network partner institute HCMR who also published all sequence data. Raw sequence files of the first data release package are available for download at the European Nucleotide Archive (ENA) (Yuan et al., 2023) through the accession numbers provided via the GitHub repository (see Supplementary Table S1 for link and Supplementary File 1 for accession numbers) and under the umbrella study PRJEB72316 (https://www.ebi.ac.uk/ena/browser/view/PRJEB72316). The accession numbers for the sequencing negative control samples are also included there. Information on the demultiplexing procedures applied on sequencing output data is provided as well, denoting if sequence reads deposited on ENA contain primer sequences or not; see Supplementary Table S1 for link and Supplementary File 1). Overall, the sequencing data comprises 200 samples (190 material samples plus ten samples which were re-sequenced due to initially low read yield) for the COI and 18S marker genes, and 111 samples for ITS, plus four to nine negative control samples per marker gene (Supplementary File 1).

### Bioinformatics processing of raw sequence data

To deposit taxonomic observations derived from the COI, 18S and ITS marker genes in the European Ocean Biogeographic Information System (EurOBIS), sequence data were processed with the Pipeline for Environmental DNA Metabarcoding Analysis, PEMA v.2.1.4 (Zafeiropoulos et al., 2020). See Supplementary Text S1 details on bioinformatics processing. Given that different parameter settings can lead to rather different outcomes (Brandt et al., 2021; Zafeiropoulos et al., 2020), a fixed set of parameters was used for each marker gene and sequence data were processed separately for each sequencing run. All parameter files used for each PEMA run are available at the dedicated repository (see Supplementary Table S1 for link). For the 18S marker gene, sequences were clustered into operational taxonomic units (OTUs) using the VSEARCH v2.9.1 algorithm (Rognes et al., 2016) with a threshold of 0.97, while for ITS and COI, clustering was performed with Swarm v2 (Mahé et al., 2015), applying a threshold of d = 10 to infer amplicon sequence variants (ASVs). Note that PEMA initially defined the result of Swarm processing as inferred ASVs, i.e., sequences which differ by one or more nucleotides, which is now corrected to swarm-clusters (Hakimzadeh et al., 2023). We here use PEMA’s initial terminology for consistency reasons. Taxonomy was assigned to 18S OTUs and ITS ASVs with the CREST LCAClassifier v3.0 (Lanzé et al., 2012), using the PR2 v.4.13.0 (Guillou et al., 2013) and Unite v7.2 (Nilsson et al., 2018) databases, respectively. For COI sequences, taxonomic annotation was performed using the RDP classifier (Wang et al., 2007) with the MIDORI database v2.0 (Machida et al., 2017). Singletons and OTUs/ASVs unclassified at domain level were removed. Abundances of OTUs/ASVs present in negative control samples were adjusted. All bioinformatics analyses were supported by the High Performance Computing system of the Institute of Marine Biology, Biotechnology and Aquaculture of HCMR (Crete, Greece) (Zafeiropoulos et al., 2021).

### Data management, EurOBIS submission, and the ARMS-MBON GitHub space

The data management proceeded in multiple stages: (i) collecting the (meta)data from the field scientists and from the processing of genetic data, (ii) harvesting and organising these in a public space, and quality controlling the metadata, (iii) adding semantics, organising the data in Research Object Crates (RO-Crates; https://www.researchobject.org/ro-crate/) (Peroni et al., 2022) and adding provenance, and (iv) submitting the data to EurOBIS. These steps were performed as follows:

(i) The types of (meta)data that were collected from the sampling teams of each observatory are outlined in Obst et al. (2020). The event metadata (observatory, event, and sample metadata, ENA accession numbers for sequencing data) were collected via a project spreadsheet (Google Sheets) and via the ARMS-MBON project space on the PlutoF platform (https://www.plutof.ut.ee), and data (ARMS plate images and spreadsheets) were uploaded by each team to PlutoF. The PEMA parameter files and selected outputs (taxonomic classifications and fasta files for sequencing data) from the bioinformatics processing were uploaded to a Google Drive folder.
(ii) The ARMS-MBON GitHub space (see Supplementary Table S1 for links) was chosen for all subsequent steps of the data management. Particular reasons for choosing GitHub were: its ease of access (within and external of the team); the ability to track changes, to implement custom workflow support through actions, and to build and host web-page-like “landing pages”. The Google sheets and data on the Google Drive were harvested into respective GitHub repositories, as was the metadata from PlutoF (as Java Script Object Notation, JSON). All harvested data were quality controlled (i.e., consistency checks, correction of mistakes) and combined into a sampling event, an image, an omics (i.e., sequencing data), and an observatory spreadsheet (comma separated-values, CSV); each data spreadsheet is accompanied by a metadata spreadsheet that defines and adds semantics to the column names. Due to their file size, the images themselves were not uploaded to GitHub, but their PlutoF (open access) URLs were, and these are included together with image metadata in a CSV file (an improved image database will be developed to better archive, annotate, and serve the ARMS-MBON image data). The PEMA input and output files were uploaded to the *processing_batch1* repository. Due to their large combined size, the fasta files were instead uploaded to the Marine Data Archive (MDA) (https://www.mda.vliz.be) and their (open access) URLs included in the relevant GitHub folder (see Supplementary Table S1 for respective links to all here described data products).
(ii) Once the data were organised into repositories in GitHub, we packed each repository as an RO-Crate. Within the RO-Crate JSON file, the contents of the repository are described and provided with provenance metadata using controlled vocabularies, thus making them machine-accessible and interoperable.
(iv) The ARMS-MBON data release package from the first sampling campaign (i.e., all taxonomically classified occurrences from the marker gene analysis with a minimum of two sequence reads) has been formatted into Darwin Core Archives (DwCA) for each marker gene for submission to EurOBIS. These data have been submitted using the relatively new DNA extension, and EurOBIS has been working on the inclusion of these data in their database, hence, these data will be published in EurOBIS in the autumn of 2024. A metadata record for these data has been created in the Integrated Marine Information System (IMIS; see Supplementary Table S1 for links) and all current and future links to the data are accessible there. The DwCAs, as well as all associated observatory and sampling event metadata, ENA accession numbers of amplicon sequencing data and links to image data specifically for the first data release package are accessible via the *data_release_001* repository on GitHub (see Supplementary Table S1 for link). The metadata files there represent subsets of the corresponding quality-controlled *combined* metadata files which contain information on all observatories, sampling events, image data and genetic data of ARMS-MBON to date (i.e., not only for the sampling events described here). The *analysis_release_001* repository on GitHub comprises all relevant bioinformatics analysis data (i.e., parameter files and outputs of PEMA processing) associated with the EurOBIS submission. This repository is merely a duplicate of the *processing_batch1* repository. All code used for exploratory data analysis described below can be found in the *code_release_001* repository. See Supplementary Table S1 for links to the repositories.

### Exploration of sequencing data

We explored the PEMA-processed sequencing data to present potential directions of utilising ARMS-MBON data sets. See Supplementary Text S2 for details. Briefly, data from individual PEMA runs we provide on GitHub were merged for each marker gene and further curated to obtain a data set for visualisations and ecological assessments. As no confidence threshold was applied within PEMA for taxonomic assignments of COI ASVs (note that this is therefore also the case for the EurOBIS submission and users are urged to apply their own self-chosen cut-off), we excluded all rank assignments with a confidence value of below 0.8. We further removed certain samples and replicates to solely assess the ARMS mobile and sessile data and to reduce diversity inflation. Potential contaminant sequences were discarded.

We determined the number and/or abundance of unique phyla, species identified, ASVs/OTUs classified to species level, and species shared between the data sets of the three marker genes. Where reference databases did not provide correct phylum level classification, we manually added these. In terms of alpha diversity, we assessed the observed ASV/OTU richness and the number of identified species across observatories, as well as frequency distributions of these two parameters (i.e., re-occurrence of ASVs/OTUs and species identified across observatories). We also determined the influence of sampling effort on diversity variables. Here, we computed Spearman’s correlation of sequencing depth and ARMS deployment duration versus ASV/OTU richness and the number of species identified in each sample. Furthermore, we computed Spearman’s correlation of the number of ARMS units deployed and the number of samples included in the analysis post-curation versus ASV/OTU richness and the number of species identified at each observatory. Where the correlation was statistically significant (*p* < 0.05) and moderate to strong (Spearman’s *ρ* > 0.5), we performed analysis of simple linear regression to model the relationship between sampling effort predictor variables and dependent diversity variables.

In order to test the application potential of the derived species observation data, we performed a scan against reference checklists for ecological key species. Species occurrences with at least two sequence reads were scanned against the following databases: i) AZTI’s Marine Biotic Index (AMBI) (Borja et al., 2000, 2019) for species very sensitive to disturbance; ii) the World Register of Introduced Marine Species (WRiMS) (Costello et al., 2021, 2024) for species with alien status at the place of observation; and iii) the Red Lists of the International Union for Conservation of Nature (IUCN) and Baltic Marine Environment Protection Commission (Helsinki Commission, HELCOM) for species registered as Near Threatened, Vulnerable, Endangered or Critically Endangered. The AMBI and IUCN/HELCOM information were obtained using the World Register of Marine Species’ (WoRMS) (Ahyong et al., 2024) REST services (https://www.marinespecies.org/rest/), while the WRiMS checks can be replicated using the Jupyter notebook on https://www.github.com/vliz-be-opsci/lw-iji-invasive-checker. Occurrences of red-listed species were confirmed by scanning against known distribution in WoRMS and corrected where necessary.

Samples were tested for differences in alpha diversity among locations with varying degrees of anthropogenic influence (i.e., industrial, semi-industrial, low human influence (LHI), and protected; see Supplementary File S1). For statistical comparison, samples with less than 5,000 reads were removed and the remaining samples rarefied to an equal sequencing depth. Samples classified as “industrial” were grouped into one category (“industrial/semi-industrial”) with samples classified as “semi-industrial”. Mean and standard deviation (SD) of the two alpha diversity measures were calculated for samples of each influence type. Data was subsequently checked for normality and log(1+x)-transformation applied where necessary. Subsequently, unidirectional analysis of variance (ANOVA) was performed to test for differences between influence types, with post-hoc Tukey’s test for pairwise comparisons. Where normality could not be achieved through transformation, non-parametric Kruskal-Wallis rank sum test was applied. All code used for exploratory analysis can be found at the dedicated GitHub repository (see Supplementary Table S1). Analyses and data visualisation were performed in R v4.1.0 (R Core Team, 2021) via RStudio v2022.07.1 (RStudio Team, 2022) using packages of the tidyverse v1.3.1 collection (Wickham et al., 2019) and the packages Biostrings v2.60.2 (Pagès et al., 2020), phyloseq v1.36.0 (McMurdie & Holmes, 2013), vegan v2.6.2 (Oksanen et al., 2023), ggpubr v0.4.0 (Kassambara, 2020), grafify v4.0 (Shenoy, 2021), plyr v1.8.7 (Wickham, 2011), scales v1.3.0 (Wickham et al., 2023), egg v0.4.5 (Auguie, 2019), UpSetR v1.4.0 (Conway et al., 2017), xlsx v0.6.5 (Dragulescu & Arendt, 2020), writexl v1.5.0 (Ooms, 2024), and openxlsx v4.2.5 (Schauberger & Walker, 2021).

## RESULTS

### Overall description of the data set

Out of the 200 sequenced sample units (i.e., 190 material samples representing biological and technical replicates, plus 10 re-sequenced samples) from the first ARMS-MBON sampling campaign, 195 and 200 were successfully sequenced (i.e., samples containing sequences that are deposited on ENA) using the COI and 18S marker genes, respectively, while 111 samples (out of 111 samples included for this gene) were successfully sequenced using the ITS marker (Table 1). Sequence processing of these samples using PEMA resulted in 54,641, 11,294 and 10,280 unique ASVs/OTUs for COI, 18S and ITS, respectively (Table 2). After further curation and filtering (i.e., negative-control-correction and removal of unclassified sequences), 189 samples with 51,782 ASVs and 9,596 OTUs remained for the COI and 18S data sets, respectively. For ITS, 42 samples with 508 ASVs remained (note that for ITS, ASVs could be inferred from two sequencing runs only with the pipeline and parameters applied here, i.e., for runs of July 2019 and April 2021). This corresponded to 1,567,301 sequence reads for COI, 3,910,167 sequence reads for 18S and 49,782 sequence reads for ITS (Table 2). All occurrence records with a minimum of two sequence reads of these classified sequences will be accessible through EurOBIS, as explained above. From the COI, 18S and ITS data sets, 18,402, 21,482, and 493 occurrences will be deposited, respectively (Table 2, see Supplementary Table S1 for links to IMIS metadata and DOI records).

**Table 2.** Overview of results from the sequence data processing using PEMA and resulting EurOBIS data sets.

|  | COI | 18S | ITS |
| --- | --- | --- | --- |
| Overall number of unique ASVs/OTUs pre-curation | 54,641 | 11,294 | 10,280 |
| Number of PEMA-processed samples with classified ASVs/OTUs remaining after negative-control-correction (excl. negative controls) | 189 | 189 | 42 |
| Overall number of unique, classified, negative-control-corrected ASVs/OTUs | 51,782 | 9,596 | 508 |
| Sequencing depth (total read number of unique, classified, negative-control-corrected ASVs/OTUs) | 1,567,301 | 3,910,167 | 49,782 |
| Number of occurrences deposited in EurOBIS | 18,402 | 21,482 | 493 |

### Taxonomic profiles of three marker gene data sets

To explore the ecological properties of the processed sequencing data, we further curated the data set as described above (i.e., applying classification confidence threshold for COI, exclusion of some samples and removal of potentially spurious sequences, etc.). This curated data set – an illustration of how a potential user-curated data set may look – comprised a subset of 162 samples for COI and 18S, respectively, and 34 samples for ITS (Table 3). These samples contained 40,363, 8,700 and 372 ASVs/OTUs with 1,223,460, 2,875,245 and 24,978 sequence reads for the COI, 18S and ITS data sets, respectively. Application of these three marker genes led to the recovery of 65 eukaryotic phyla, of which 38, 57 and 9 were present in the COI (at the confidence threshold of 0.8 applied here), 18S and ITS data, respectively (Table 3, see Supplementary File 2 for further details). In regards to relative read abundance, the COI data set classified to phylum level was dominated by metazoan phyla (i.e., eight out of the ten most abundant phyla) – mainly Arthropoda, Annelida, Chordata, Bryozoa and Cnidaria – while almost half of all reads (∼44%) belonged to sequences unclassified at phylum level (Figure 2A, Table 3 and Supplementary File 2). For 18S, metazoans of the phyla Chordata, Mollusca, Arthropoda, Annelida and Cnidaria dominated the data set, but more non-metazoan phyla were among the most abundant taxa compared to the COI data (Figure 2C and Supplementary File 2). The majority of 18S reads could be classified to at least phylum level (∼97%; Table 3). As expected, mainly fungal phyla were recovered using the ITS marker gene (Cnidaria was the only non-fungal phylum in the ITS data) and Ascomycota made up more than half of all reads in this gene’s data set (Figure 2E and Supplementary File 2). Approximately two-thirds (∼61%) of ITS reads were classified to the phylum level (Table 3).

**Figure 2.**
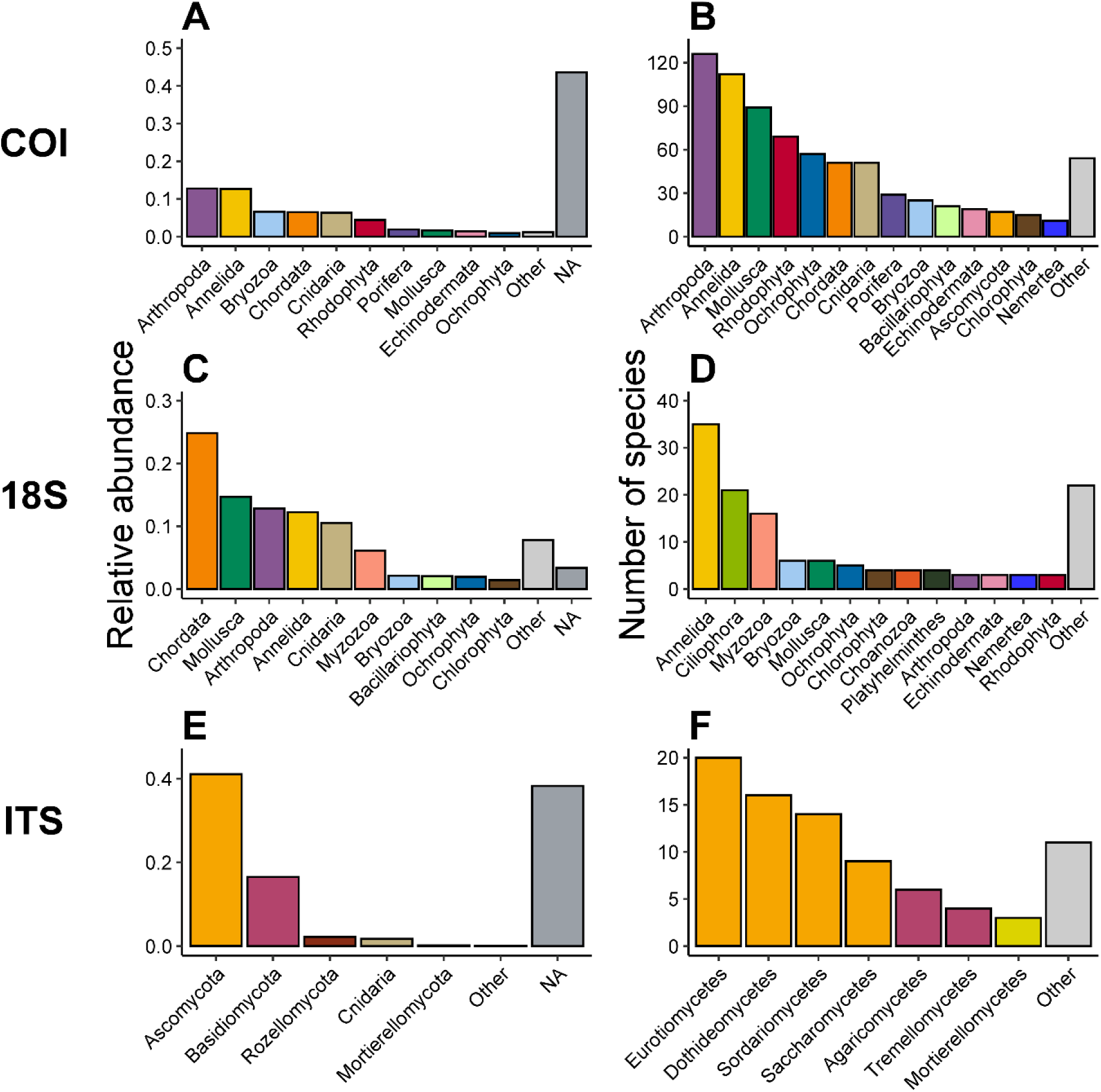
Relative read abundance of the ten most abundant phyla in the COI **(A)** and 18S **(C)** and five most abundant phyla in the ITS data sets **(E)**. Less abundant phyla are grouped as *Other*, while relative abundance of sequence reads unclassified at phylum level are grouped as *NA*. Number of identified species within each phylum for COI **(B)** and 18S **(D)** and within each class for ITS **(F)**. Phyla / classes with less than ten (i.e., COI) or three (i.e., 18S and ITS) identified species are grouped as *Other*. Colours correspond to the same unique phyla across all plots. Class level representation was chosen in **(F)** for better taxonomic resolution and colours correspond to the fungal phylum each class belongs to.

**Table 3.** Overview of curated sequence data used for ecological analysis.

|  | **COI** | **18S** | **ITS** |
| --- | --- | --- | --- |
| Number of sequencing samples | 162 | 162 | 34 |
| Overall number of ASVs/OTUs | 40,363 | 8,700 | 372 |
| Sequencing effort (total read number) | 1,223,460 | 2,875,245 | 24,978 |
| Number of phyla recovered | 38 | 57 | 9 |
| Percentage of reads classified at phylum level (with the confidence thresholds applied here) | 56.39 | 96.65 | 61.15 |
| ASV/OTUs with Linnean species name classification (with the confidence thresholds applied here) | 2,220 | 399 | 92 |
| Number of unique species identified with Linnean name | 746 | 135 | 82 |
| Derived species observation records (occurrences with a minimum of two sequence reads in a sample) | 2,772 | 984 | 106 |
| Number of phyla represented in species identifications | 35 | 31 | 4<br>(15 classes) |

The use of the COI marker gene resulted in the most species identifications (at the confidence threshold of 0.8 applied here) and observations (occurrences with more than one read in a sample) compared to the use of 18S and ITS. We recovered 2,220 ASVs with species level classification from the COI data, corresponding to 746 unique species and 2,772 observation records (i.e., occurrences of at least two reads; Table 3). For 18S and ITS, 399 and 92 OTUs/ASVs could be assigned a species name, which represented 135 and 82 unique species with 984 and 106 species observations, respectively (Table 3). In total, the species identifications represented 45 unique eukaryotic phyla: 35 for COI, 31 for 18S and four fungal phyla for ITS (Table 3, see Supplementary File 2 for more details). In the case of COI, more than 100 species were identified for both the Arthropoda and Annelida phyla, with Molusca, Rhodophyta, Ochrophyta, Chordata and Cnidaria also representing a large share of the species level identifications (Figure 2B and Supplementary File 2). Contrarily, Annelida, Ciliophora and Myzozoa dominated species occurrences in the 18S data set (Figure 2D and Supplementary File 2). Fungal species identifications retrieved from the ITS data mainly belonged to classed of Ascomycota (i.e., Eurotiomycetes, Dothideomycetes, Sordariomycetes, etc.) and Basidiomycota (i.e., Agaricomycetes, Tremellomycetes, etc.) (Figure 2F and Supplementary File 2). The three marker genes recovered relatively distinct groups of species, as only one (i.e., ITS vs. 18S and ITS vs. COI) and 18 (i.e., COI vs. 18S) of the species recovered here overlapped between the three data sets (Supplementary Figure S1), and 727, 116 and 80 identified species were unique to the COI, 18S and ITS data sets, respectively. No species occurred in all three marker gene data sets (Supplementary Figure S1).

### Genetic and species diversity at ARMS-MBON observatories

Alpha diversity of genetic units (i.e., ASVs and OTUs) and identified species varied between observatories. Between 60 (Eilat - Israel) and 246 species (Koster - Sweden) from COI data (at the confidence threshold applied here) and 2 (i.e., Getxo - Spain) and 53 species (Koster - Sweden) from 18S data were identified at the observatories (Figure 3; Supplementary Table S2). Richness of ASVs/OTUs ranged between 365 (Læsø - Denmark) and 5,472 (TZS - Finland) for COI and between 253 (Getxo - Spain) and 2,136 (Koster - Sweden) for 18S (Figure 3; Supplementary Table S2). While the high 18S OTU diversity at Koster on the Swedish west coast aligned with its high number of identified species, the high COI ASV richness at the TZS observatory on the Gulf of Finland contrasted its relatively low number of identified species (n=70) compared to observatories with lower genetic diversity. In comparison, the two ARMS from the Norwegian coast at Bodø captured more than twice as many species (n=168) as the TZS observatory but showed less than half the genetic diversity (2,275 ASVs) for COI. It should be noted that alpha diversity was driven by differences in sampling effort and sequencing depth to some extent (see below).

**Figure 3.**
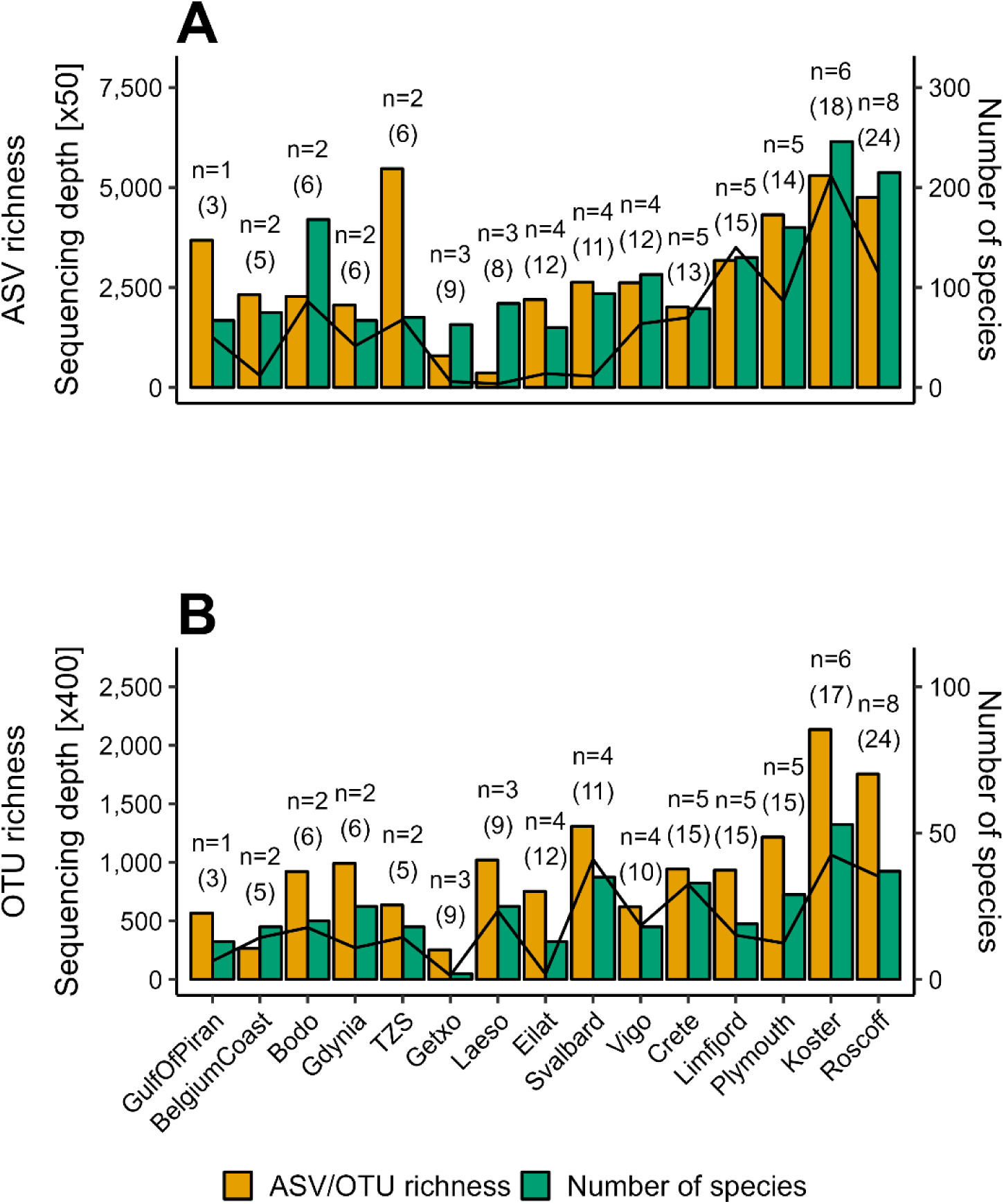
Diversity observed for the COI **(A)** and 18S **(B)** data sets across the 15 observatories measured as ASV/OTU richness (yellow; shared axes on the left) and the number of identified species (green; axes on the right) at the classification confidence threshold applied here. The number of ARMS units deployed at the respective observatory is shown on top of the bars (n). Numbers in parentheses equal the number of genetic samples remaining for each observatory after curation and filtering for ecological analysis. The black line denotes sequencing depth (i.e., cumulative sequence read number of all genetic samples for each observatory; left shared axes). Observatories are ordered from left to right by an increasing number of ARMS units deployed.

Analysis of frequency distributions across observatories showed that ∼94% (n = 38,090) and ∼4% (n = 1,593) of COI ASVs occurred at one or two observatories only, respectively (Figure 4A). Frequency distribution was less skewed for 18S data, with ∼67% (n = 5,794) and ∼18% (n = 1,565) of OTUs found at one or two observatories only, respectively (Figure 4B). Approximately 400 species (∼53%) identified from the COI data occurred at one observatory only, while 160 species (∼21%) occurred at two observatories (Figure 4C). A relatively low number of species identified from the COI data occurred at 10 or more observatories (n = 13, ∼2%; Figure 4C). Half of all species (n = 68) identified using the 18S marker gene were found at one observatory only, and 17 species (∼13%) occurred at only two observatories (Figure 4D). Two species identified from 18S data appeared at ten or more observatories (Figure 4D).

**Figure 4.**
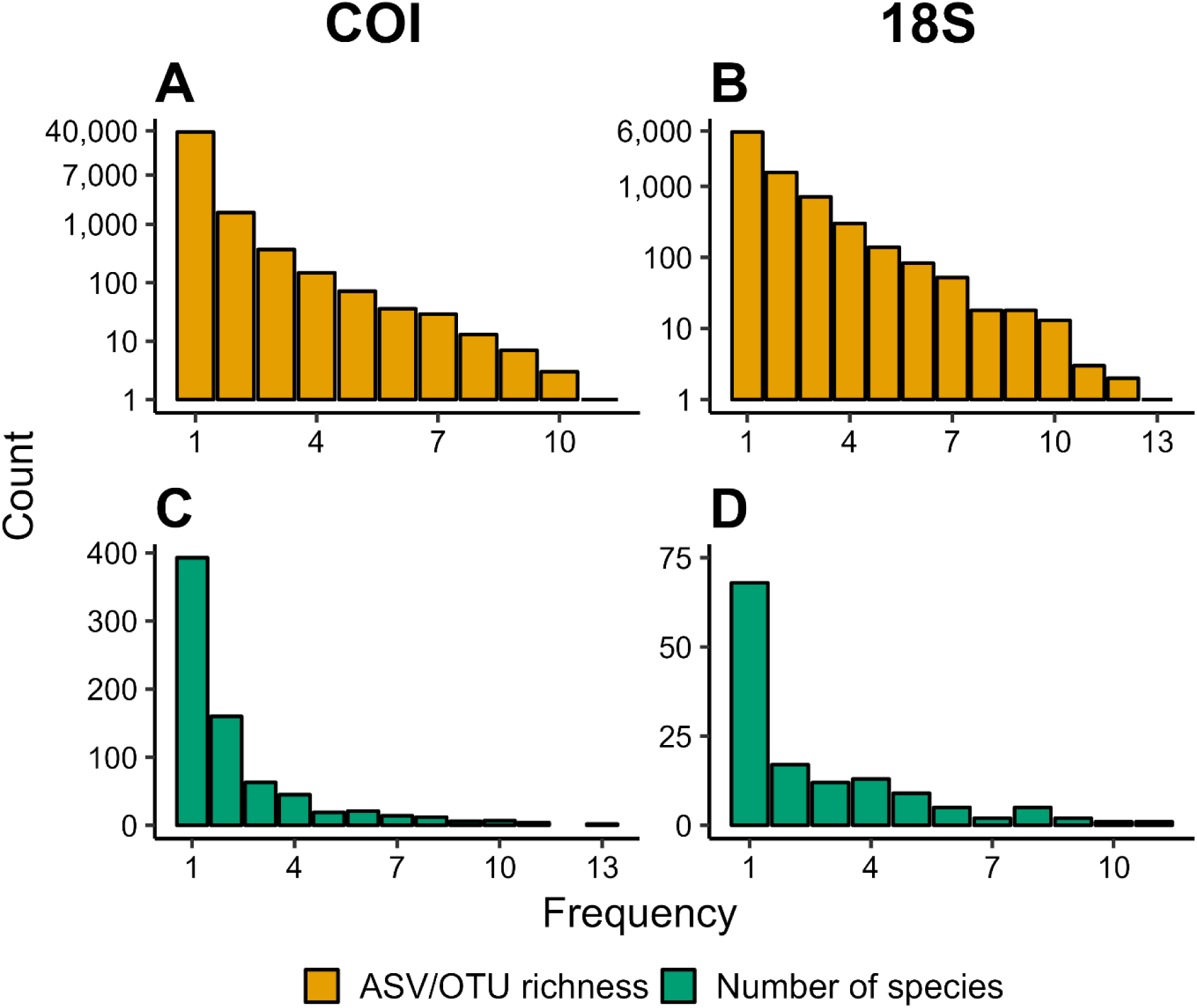
Frequency distribution of ASVs for COI **(A)** and OTUs for 18S **(B)** and of species identified for COI **(C)** and 18S data **(D)**. Counts represent the number of entities (i.e., ASVs/OTUs or species identified) occurring at a given frequency, in this case the number of observatories. Note the log_2_-transformation of the y-axis in **A** and **B**.

### Influence of sampling effort and deployment duration on diversity measures

During the first ARMS-MBON sampling campaign, observatories deployed one (GulfOfPiran - Slovenia) to eight (Roscoff - France) ARMS units, for periods ranging from 37 (BelgiumCoast - Belgium; UnitID AZFPin) to 649 days (Eilat - Israel; UnitID Katza2) (Figure 1B; see Supplementary File 1 for further details). After data curation and filtering, this resulted in a data set of three (GulfOfPiran - Slovenia) to 24 samples (Roscoff - France) with at least one sequence read for each observatory for both the COI and 18S marker genes (Figure 3). Given differences in sequencing depth and quality, not all of the three fraction samples remained for each ARMS unit post-curation.

Computation of correlation indicated no significant linear association between deployment duration and the number of species identified in each sample for both marker genes (Figure 5A, B; see Supplementary Table S3 for detailed results of correlation analysis). This was also the case for deployment duration vs. ASV/OTU richness (Figure 5A, B). For both marker genes, sequencing depth (i.e., the number of sequence reads) significantly drove the number of species identified and the ASV/OTU richness observed in each sample, with moderate to strong correlation (*p* < 0.001 and Spearman’s *ρ*(160) ≥ 0.60 for all associations; Figure 5C, D).

**Figure 5.**
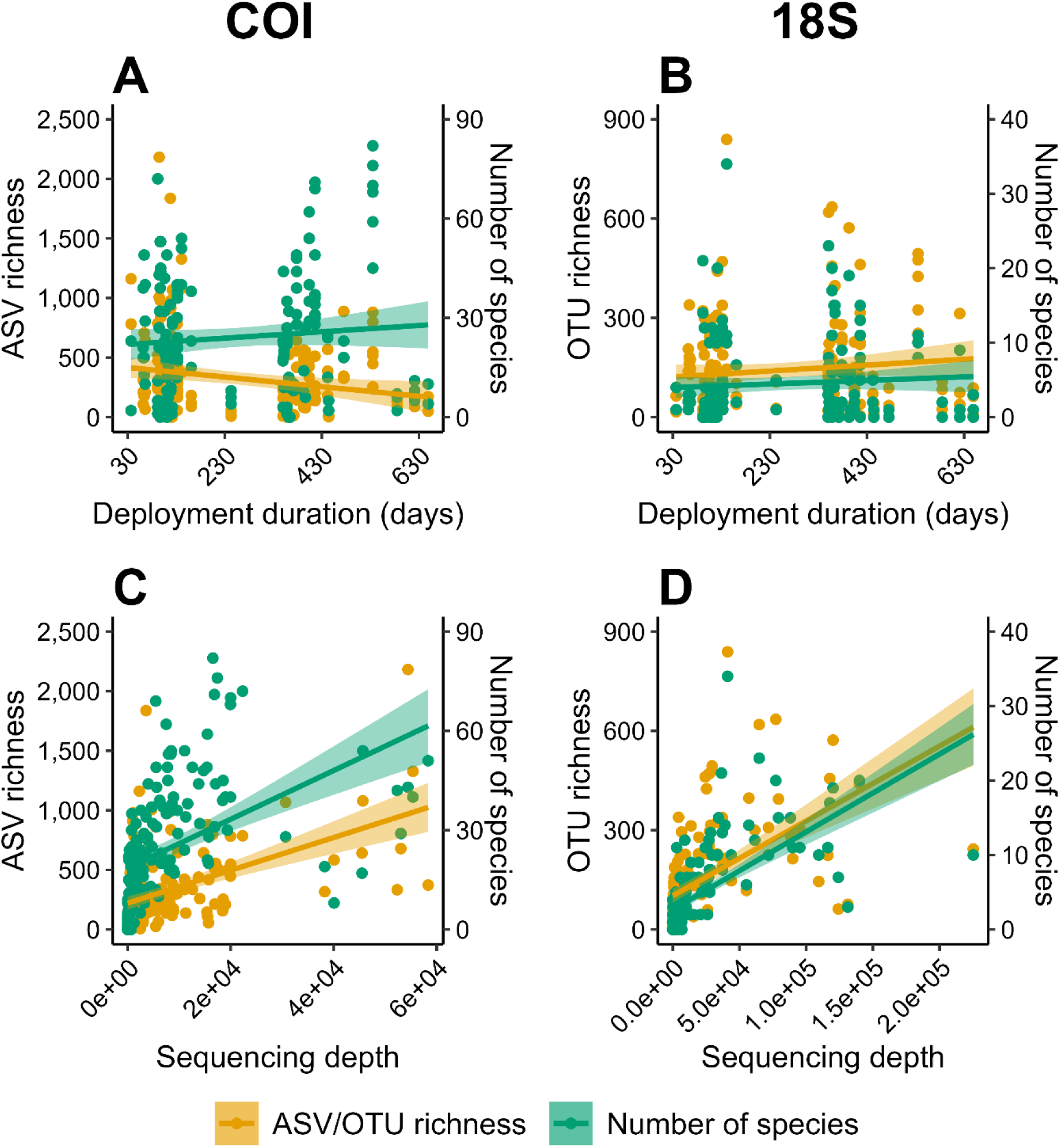
Relationship of the observed ASV/OTU richness (yellow; axes on the left) and the number of species identified (green; axes on the right) in each sample versus the deployment duration in days of the respective ARMS units for the COI **(A)** and 18S **(B)** data sets; and relationship of ASV/OTU richness (yellow; axes on the left) and the number of species identified (green; axes on the right) in each sample versus sequencing depth (i.e., number of sequence reads in each sample) for the COI **(C)** and 18S **(D)** data sets. Solid lines represent linear regression for ASV/OTU richness (yellow) and the number of species identified (green), shaded areas represent the corresponding 95% confidence intervals. No significant linear correlation was found for associations in **A** and **B**.

Analysis of linear regression indicated that sequencing depth accounted for around 23% to 41% of variation in ASV/OTU richness and the number of species identified in each sample for both marker genes. However, residual standard errors suggested the regression models did not fit the data accurately, although fit was more accurate for 18S compared to COI data (see Supplementary Table S4 for detailed results of regression analysis).

For the COI marker gene, only the number of species identified at each observatory was significantly and moderately to strongly correlated to both the number of ARMS units deployed and the number of samples included in the analysis (*p* = 0.018, Spearman’s *ρ*(13) = 0.60; and *p* = 0.021, Spearman’s *ρ*(13) ≥ 0.59, respectively; see Supplementary Figure S2A, C and Supplementary Table S3). The observed COI ASV richness did not show a significant linear relationship with these two sampling effort parameters (Figure S2A, C and Supplementary Table S3). In the case of 18S, both the observed OTU richness and the number of species identified at each observatory significantly and moderately correlated with both the number of ARMS units deployed and the resulting number of samples included in the analysis (*p* ≤ 0.012 and Spearman’s *ρ*(13) ≥ 0.64 for all associations; see Supplementary Figure S2B,D and Supplementary Table S3). Analysis of linear regression indicated that the number of ARMS units deployed and the number of samples included accounted for around 37% to 51% of variation in the number of species identified at each observatory for both marker genes. In the case of 18S, the number of ARMS units deployed and the number of samples included were both responsible for ∼52% of variation in the OTU richness observed at the observatories. Given relatively high variation in the dependent variables and therefore considerable residual standard errors, observatory-wise regression models did not fit our data accurately as described above for sample-wise models (Supplementary Table S4).

### Identification of ecological key species

The scan against databases for sensitive, non-indigenous, and red-listed species resulted in observations of species in all three categories across the observatories (Figure 6). Overall, we observed 88 species registered in AMBI as sensitive to disturbance, 32 species listed as “alien” in WRiMS at the location of occurrence, and 4 species registered as near threatened, vulnerable, endangered or critically endangered across all observatories (Supplementary Table S5 and Supplementary File 3). The observatory at Koster (Sweden) detected the highest number of sensitive species (n = 37), while Limfjord (Denmark) displayed the highest number of non-indigenous species (NIS). Red-listed species were only detected at the Plymouth (UK) (*Mya truncata*), Koster (Sweden) (*Cliona celata* and *Echinus esculentus*) and Roscoff (France) (*Cliona celata* and *Nucula nucleus*) observatories.

**Figure 6.**
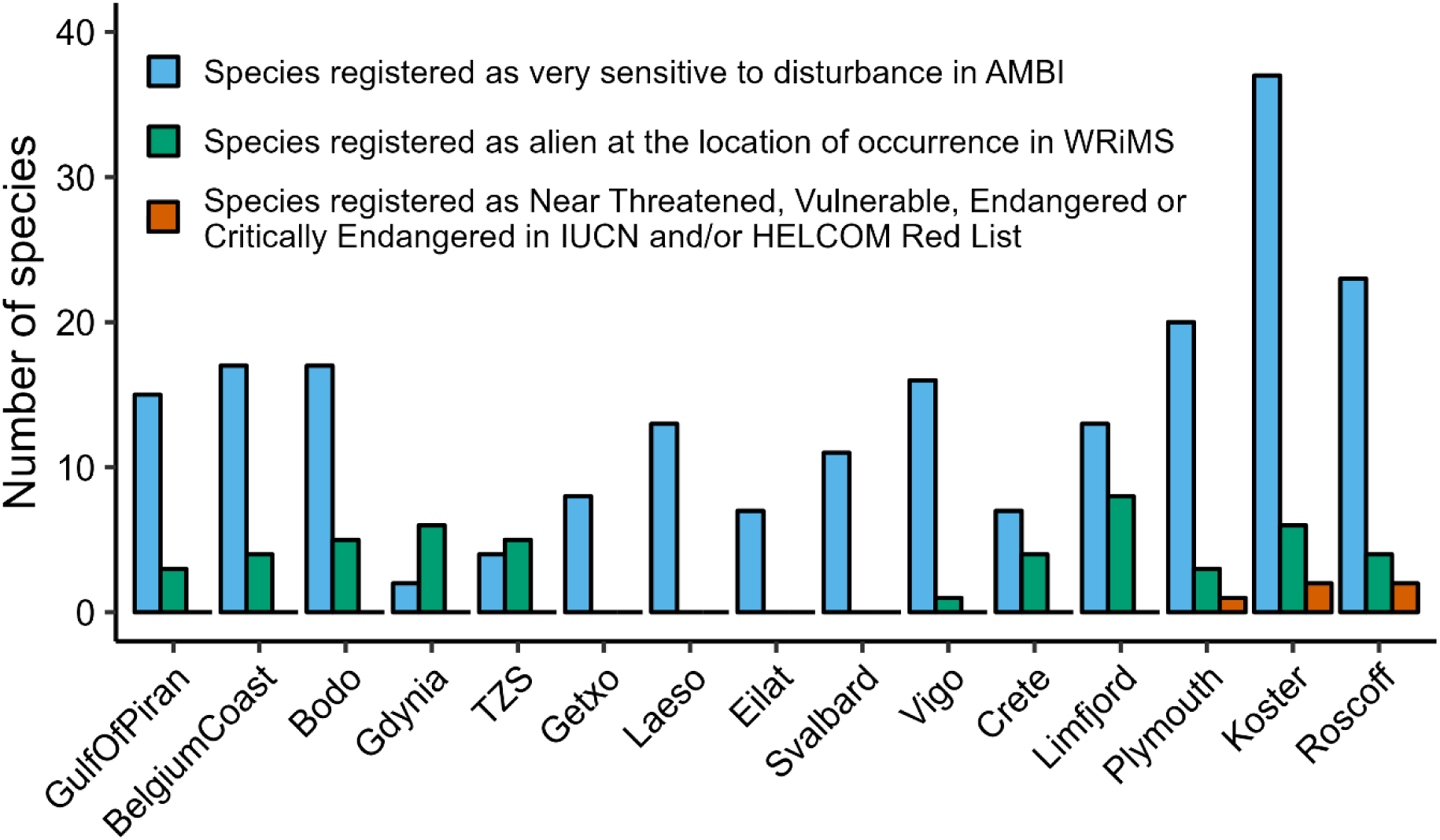
Number of identified species (with occurrences of at least two sequence reads) across observatories listed in four different databases. Data of COI and 18S marker genes were pooled for each observatory. Blue – species registered as very sensitive to disturbance in AMBI; green – species registered as alien at the location of occurrence in WRiMS; orange – species registered as Near Threatened, Vulnerable, Endangered or Critically Endangered in the Red Lists of the IUCN and HELCOM. Observatories are ordered from left to right by increasing number of ARMS units deployed (n = 1 for GulfOfPiran, n = 8 for Roscoff).

### Alpha diversity across locations with varying degrees of anthropogenic influence

The number of identified species differed significantly among habitats with varying degrees of anthropogenic influence based on COI data (low human influence: 26 ± 8, industrial/semi-industrial: 29 ± 13, protected: 38 ± 15; ANOVA: *F*_2,60_ = 5.043; *p* = 0.009) (Figure 7A and Supplementary Table S6) but based on 18S data they did not (industrial/semi-industrial: 5 ± 5, low human influence: 6 ± 4, protected: 7 ± 3; Kruskal-Wallis χ^2^ = 2.543, *p* = 0.280) (Figure 7B and Supplementary Table S6). Mean ASV richness ranged from 216 ± 90 (low human influence) to 260 ± 200 (industrial/semi-industrial) and 260 ± 173 (protected) for COI data (Figure 7A) and no statistically significant difference was detected among habitat types given high within-group variability (ANOVA of log(1+x)-transformed data: *F*_2,60_ = 0.241; *p* = 0.787; see Supplementary Table S6 for details of statistical tests). The same was observed for 18S data (ANOVA: *F*_2,60_ = 1.906; *p* = 0.157), with OTU richness ranging from 123 ± 78 (industrial/semi-industrial) to 136 ± 53 (low human influence) and 165 ± 84 (protected) (Figure 7B and Supplementary Table S6).

**Figure 7.**
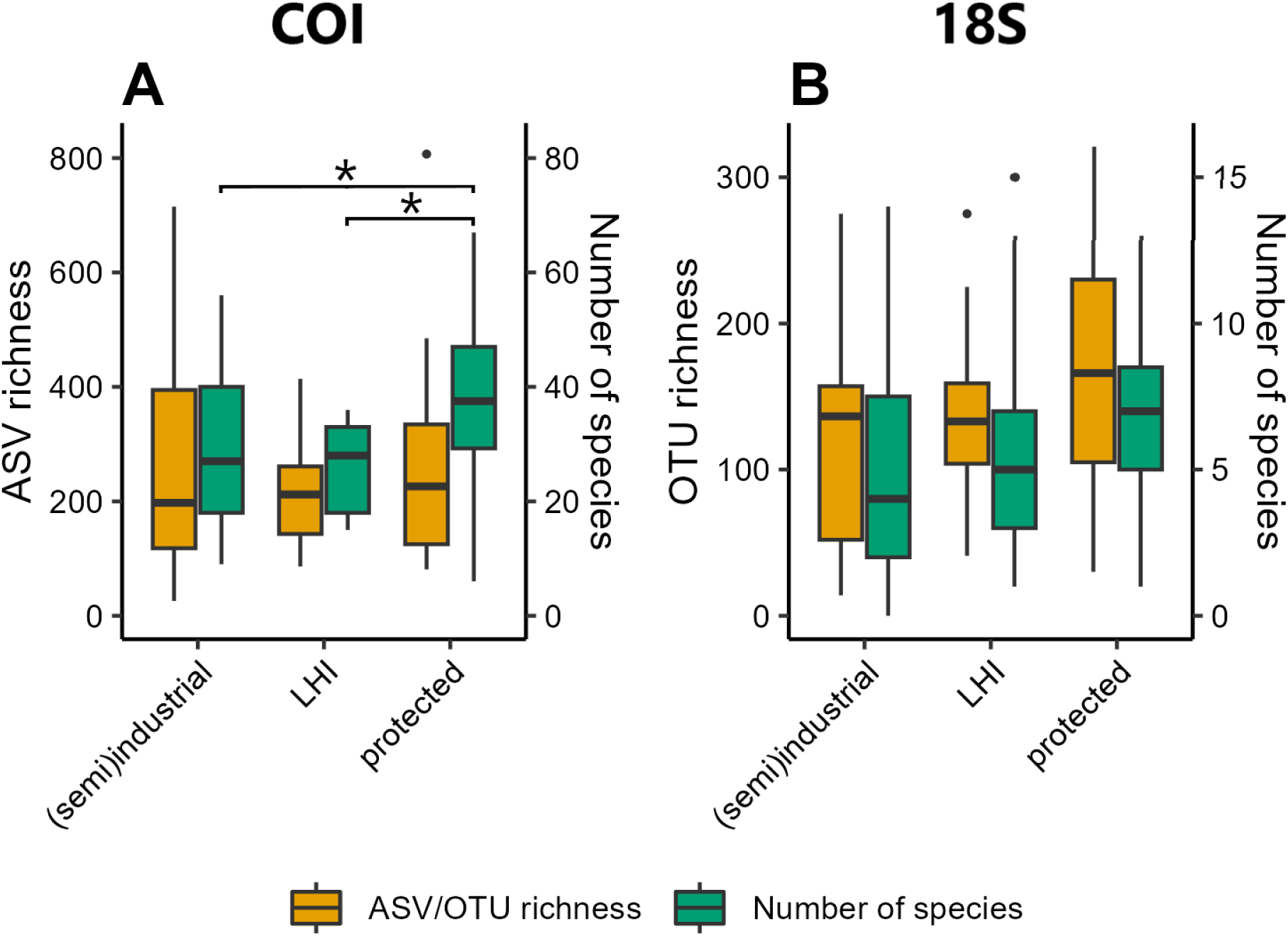
Alpha diversity across locations with varying degree of anthropogenic influence (i.e., industrial/semi-industrial, LHI - low human impact, and marine protected area). Boxplots show alpha diversity measured as ASV/OTU richness (yellow; axes on the left) and the number of identified species (green; axes on the right) at the classification confidence threshold applied here for samples of each influence type for the COI **(A)** and 18S **(B)** data sets. Boxes span the interquartile range (IQR) from first to third quartile. Horizontal lines within boxes represent median values. Whiskers extend to minimum (lower) and maximum (upper) values up to 1.5*IQR beyond either side of the IQR. Values falling outside this range are outliers represented by single black dots. Significant differences in the number of identified species for the COI data set according to ANOVA are shown in **(A)** by horizontal bars with asterisks above boxplots (*: *p* < 0.05).

## DISCUSSION

Here we present the data set of raw and processed amplicon sequencing data from the first sampling campaign (2018-2020) of ARMS-MBON. To the best of our knowledge, this campaign represents the most geographically widespread sampling initiative using ARMS to date. Raw sequencing (and image) data are open-access and, thus, they can be subjected to re-processing by all interested users according to their needs. In addition, we processed sequencing data using a dedicated pipeline, and all taxonomic occurrences will be published on EurOBIS. This processed data set was further filtered and curated, representing a potential use case for ecological analysis or taxonomic screening. Analysis of this curated data set shows that ARMS are able to capture eukaryotic taxa belonging to more than 60 phyla, while composition of the hard-bottom benthic cryptofauna in terms of dominant phyla was comparable with those of earlier studies using ARMS (e.g., Carvalho et al., 2019; Ip et al., 2022; Pearman et al., 2018, 2020). Our data set also allows for comparisons of communities at all taxonomic levels (i.e., including genus, class, order level identifications) which are typically applied in local and regional community studies (Staehr et al., 2022). Although we do not present any prokaryotic data here, the preserved physical samples of ARMS-MBON sampling campaigns can also be used for comparative studies of non-eukaryotic microbial communities in the future (Ip et al., 2022; Pearman et al., 2019). Finally, the catalogue of ARMS plate images collected can become a valuable data resource in the future as it will allow for analysis of benthic community composition and growth dynamics with application of advanced image classification methods (Beijbom et al., 2015).

### Recovered taxa are distinct across marker genes and observatories

The marginal overlap in identified species between the marker genes underlines the importance of applying multi-marker assays to increase taxonomic coverage in biomonitoring projects (e.g., da Silva et al., 2019; Duarte et al., 2023; Gibson et al., 2014; Gielings et al., 2021). The geographic distribution of genotypes and species identified across the ARMS-MBON network was relatively restricted with most observations of species and genotypes being unique to one or two observatories. Such pattern of sequences and taxa being unique to sample units or locations has been observed in many metabarcoding studies before (e.g., Carvalho et al., 2019; Villalobos et al., 2022) and in our case may be due to the still limited number of samples and observatories given the large geographic scale of the project. As samples and data accumulate over the coming years, it is likely that the partition with re-occurring observations will increase. In addition, the growing number of reference sequences (Porter & Hajibabaei, 2018) and/or using customised reference databases (Mugnai et al., 2023) will likewise increase the taxonomic resolution and observation records derived from the samples.

### The need to account for differences in sampling effort

Deployment periods of ARMS units ranged from 37 to 649 days, with the majority being deployed for around three months to approximately one year. Our tests showed that in most cases deployment duration did neither have a significant effect on observed species nor on genetic diversity. The weak negative correlation between deployment duration and sequence diversity in the COI data set can most likely be attributed to the low sequencing depth in some samples of ARMS deployed for longer periods (i.e., units of observatories from Svalbard - Norway and Eilat - Israel and one unit of Crete - Greece). However, we want to stress the fact that a conclusion on the effect of deployment duration can ultimately only be achieved with dedicated testing, i.e., by comparing locations with both short- and long-term deployments and equal sequencing depth. Previous studies have found contrasting results regarding the influence of deployment duration of sampling units. Using artificial substrate units (ASUs), Cahill et al. (2018) did not observe a significant effect of deployment duration on the number of specimens recovered and argued that overall recruitment patterns are predominantly driven by ecological and biogeographic conditions. In their comparative study, Leite et al. (2023) deployed ARMS and artificial seaweed monitoring systems (ASMS) for six, nine, and twelve months and found that community composition changed over time given ecological succession. The authors indicate that maximum diversity can be recovered with sampling units deployed for less than twelve months, which had also been shown by earlier studies (Leite et al., 2021). In contrast to deployment duration, we showed that recovery of sequence and taxonomic diversity significantly depend on sequencing depth. This is a well-known problem in microbial amplicon sequencing (Cameron et al., 2021; McMurdie & Holmes, 2014) and has also been shown in eukaryotic DNA metabarcoding studies (Alberdi et al., 2018; Grey et al., 2018; Shirazi et al., 2021). As expected, the number of ARMS units deployed and the number of samples analysed for each observatory significantly drove the recovered alpha diversity, as well. Large variation in ASV richness across COI samples is likely the reason that the effect of sampling effort was not significant for this specific test case. Differences in sampling effort are known to drive observed alpha diversity measures in traditional ecological surveys (Gotelli & Colwell, 2011) and, in particular, in metabarcoding studies (Evans et al., 2017; Grey et al., 2018). Hence, analytical tools need to be applied to account for them.

Given that deployment duration did not influence alpha diversity, we suggest deploying ARMS for at least three to six months during the season of most substantial growth (i.e., during spring to summer/fall or summer to fall on the respective hemisphere) to capture a representative snapshot of benthic communities and to use human, time, and material resources as efficiently as possible. We also urge for deployments of at least three ARMS units per site (i.e., with a distance of around 10 m from each other) as biological replicates to obtain a comprehensive representation of surrounding communities and to improve statistical power for comparative analysis. Given differences in sampling effort and in sequencing depth across samples – the latter being inherent to any amplicon sequencing study – we recommend that users apply statistical tools when using ARMS-MBON data to account for those during ecological analysis.

### ARMS data enable the study of large-scale biodiversity patterns

We found indications that alpha diversity of the hard-bottom benthos within marine protected areas (MPAs) is higher than in locations with more intense anthropogenic influence. Mean sequence and species richness was highest in samples from ARMS deployed in protected areas for both marker genes, but given high within-group variation across samples, this was only statistically significant for species richness of one marker gene. This can be an indication of the positive effect of protection measures (Edgar, 2011; Edgar et al., 2014) but may also be related to the fact that MPAs are typically established in areas with already low anthropogenic influence where political implementation costs are low (Devillers et al., 2015; Stevenson et al., 2020). The fact that we found higher species diversity but not higher genetic diversity in MPAs remains unexplained at this point. A possible reason for this pattern could be that high genetic diversity reflects recruitment which may be high even in habitats with anthropogenic pressures, while MPAs support species beyond the initial recruitment stage. Further, propagules and larvae of opportunistic taxa, including NIS, may be particularly ubiquitous in harbours and marinas, but less so in stable ecosystems with high conservation status such as MPAs. In addition, small, heterogeneous but isolated MPAs may support high species but low genetic diversity due to reduced connectivity (Bell & Okamura, 2005) and seascape and spatial factors may have a higher influence on genetic diversity than protection status (Benestan et al., 2023). Given the continuous sampling on a large spatio-temporal scale, data from the forthcoming sampling campaigns of ARMS-MBON can help unravel differences in diversity more clearly. In any case, ARMS data can be used to document biodiversity trends in benthic habitats in relation to human activities and eventually contribute to several essential biodiversity variables (EBVs) in the future (Kissling et al., 2018). Genetic and taxonomic data from ARMS-MBON may also be combined with open-access remote sensing data to link diversity patterns and taxonomic occurrences to environmental parameters (Pearman et al., 2019, 2020).

### ARMS are sensitive to indicator, non-indigenous, and threatened species

Our results show that ARMS can detect ecological indicator species sensitive to disturbance, which are typically used in environmental impact assessments (EIAs) (Bustos-Baez & Frid, 2003; Dauvin, 2005), environmental risk assessments (ERAs) (Kaikkonen et al., 2018), and national / regional monitoring programmes (e.g., HELCOM, 2013) to assess the health status of ecosystems. ARMS-MBON data are likely to improve such assessments since they provide information on species as well as genetic diversity, which can be analysed in relation to anthropogenic pressures. As such, ARMS should be deployed continuously in sites such as ports, marinas, wind farms, or aquaculture facilities in order to assess impacts of human activities on marine biodiversity (Witalis et al., 2021). We also detected a (low) number of red-listed species, although the fact that this was the case only for observatories with the highest sampling effort underlines the rarity of such taxa and the need for continuous and considerable sampling to track and monitor them.

The data presented here can also be used for tracking the distribution and range shift of NIS as well as other taxa (Martaeng et al., 2023; Wesselmann et al., 2024) and may be applied in various monitoring programmes and directives, such as European Union’s Marine Strategy Framework Directive (MSFD) when assessing descriptors D1 on biological diversity and D2 on NIS (Bourlat et al., 2013), or the Water Framework Directive (WFD) (Duarte et al., 2023). Such investigations may be particularly enhanced by the intraspecific diversity that can be detected through large-scale genetic data sets such as the one of ARMS-MBON, which will allow for studies of population structure or connectivity of particular species with recently proposed metaphylogeographic analyses (Antich et al., 2023; Martaeng et al., 2023; Turon et al., 2020). In addition, ARMS-MBON data are likewise useful for effective alien species matches between ports as part of same risk area assessments under the International Maritime Organization’s Ballast Water Management Convention (Stuer-Lauridsen et al., 2018). For the detection of NIS we relied on species listed as alien in WRiMS; however, this may be enhanced in the future by comparing occurrences in ARMS-MBON data sets with information in repositories such as the Global Biodiversity Information Facility (GBIF) to identify potential novel invasions or range extensions.

### How to build a successful, data-producing, long-term genetic observatory network

Lessons learned from running the ARMS-MBON project are the following:

- Standardisation: it is vital to ensure standard protocols (collecting and processing of material, managing the data) are published, understood, and followed so that the resulting data is comparable over the space and time of the observatories.
- Constant engagement: it is necessary to engage the hearts and minds of the participants for the exciting (e.g., field work) as well as the challenging and tedious (e.g., data management) parts, as otherwise the resulting (meta)data are insufficient to fulfil the project’s potential.
- Strength in numbers: it is necessary to have multidisciplinary core of experts taking control of the different parts of the project (e.g., sampling, sequencing, bioinformatics, data management, analysis), as no one scientist can do all of these with the quality and timeliness to allow for regular and trustworthy data releases.

## CONCLUSION

The ARMS-MBON initiative is a network of long-term ecological research (LTER) sites committed to the scientific exploration of hard-bottom benthic communities along the coasts of Europe and adjacent regions. Data resulting from the network’s consecutive sampling campaigns will enable the study of coastal biodiversity over large temporal and spatial scales and may crucially enhance efforts to monitor marine habitats with varying degrees of anthropogenic influences. Such long-term sampling programmes also enable improved early-detection and monitoring of specific groups of taxa, such as NIS. The strength of ARMS-MBON lies in its continuous application of standardised protocols and operating procedures, as well as centralised molecular sample processing and sequencing. These measures help reduce biases potentially introduced due to the large-scale experimental set up. As improvements to protocols and standard procedures are constantly underway, these will further enhance standardisation within the network. Importantly, the possibility of utilising either the raw data or the data processed with a standardised bioinformatics pipeline gives users the freedom to choose the data product best suited to their specific needs. As ARMS-MBON continues its sampling efforts through EMO BON, the open-access data it delivers provide an increasingly critical source of information in times of utmost urgency for large-scale environmental monitoring.

## ACKNOWLEDGEMENTS

The ARMS-MBON network was established under the infrastructure program ASSEMBLE Plus (grant no. 730984) and is currently financed by the European Marine Biological Resource Centre (EMBRC) under the program European Marine Omics Biodiversity Observation Network (EMO BON). Data publication and analysis is funded by the projects DTO-bioflow (grant agreement no. 101112823) and MARCO BOLO (grant agreement no. 101082021). Funding for individual ARMS observatories was provided by the INTERREG project GEANS (North Sea Program of the European Regional Development Fund of the European Union), the Swedish Agency for Marine and Water Management (grant no. 3181-2019), the Flanders LifeWatch contribution (Research Foundation Flanders grant I000819N), and the Aquanis 2.0 project (FONDATION Total). Data management and computational resources were provided by the Swedish Biodiversity Data Infrastructure (grant no. 2019-00242) and by IMBBC (Institute of Marine Biology, Biotechnology and Aquaculture) of the HCMR (Hellenic Centre of Marine Research). Funding for establishing the IMBBC HPC has been received by the MARBIGEN (EU Regpot) project, LifeWatchGreece RI and the CMBR (Center for the study and sustainable exploitation of Marine Biological Resources) RI. Guiding documents to obtain ABS clearance for access to genetic resources were developed under the projects INTERREG EBB (EAPA_501/2016) and H2020 EOSC-Life (grant no. 824087). JT was supported by the Carlsberg Foundation (case no. CF21-0564) and the Aage V. Jensen Foundation.

## DATA ACESSIBILITY AND BENEFIT-SHARING

### Data Accessibility Statement

All data presented in this manuscript are publicly available, see main text and Supplementary Material for detailed descriptions. Standard operating procedures and protocols are available on the dedicated ARMS-MBON GitHub repository (https://github.com/arms-mbon/documentation). All metadata and access to all image data generated by ARMS-MBON to date can be found on GitHub (https://github.com/arms-mbon/data_workspace). All genetic raw data generated by ARMS-MBON to date can be accessed on the European Nucleotide Archive (ENA) through the accession numbers provided via the GitHub repository (https://github.com/arms-mbon/data_workspace) and under the umbrella study PRJEB72316 (https://www.ebi.ac.uk/ena/browser/view/PRJEB72316). Metadata, access to image data and accession numbers for genetic data specifically for the data set presented in this manuscript are provided in the Supplementary Files and are also available on the respective GitHub repository (https://github.com/arms-mbon/data_release_001). Files used for EurOBIS submissions will be available on https://github.com/arms-mbon/data_release_001). Output files of the PEMA pipeline used as basis for EurOBIS submissions and for exploratory analysis as shown in this manuscript are accessible on https://github.com/arms-mbon/analysis_release_001. The URLs to IMIS metadata records of EurOBIS submissions and the respective DOIs will be made available by the end of April 2024. All code and input files used for exploratory analysis presented in this manuscript are available or linked to on https://github.com/arms-mbon/code_release_001.

### Benefit-Sharing Statement

Benefits Generated: ARMS-MBON represents a large-scale research collaboration with scientists from across Europe and beyond. All network partners of the observatories mentioned in this manuscript provided genetic samples and are included as co-authors. All continuously generated raw and processed data from this network is shared with the public and scientific community (see above). Our research addresses the urgent need for large-scale and long-term monitoring of marine biotic communities through extensive collaborative efforts.

## AUTHOR CONTRIBUTIONS

All authors maintained the observatories, organised the sampling campaigns, and contributed to the development of the presented methods and standards. MO, CP, KE, IS established and maintained the network. IS and CP organised the central sample processing and sequencing. ND, CP, HZ did the majority of the analytical work. KE, CP, JP, ND performed the majority of the data management. JT and PS contributed to image data management. ND wrote the final manuscript with major contributions from MO, CP, KE, JP. All other authors revised the manuscript.

## SUPPORTING INFORMATION

**Supporting Information for online publication.docx**

Contains Supplementary Texts S1 – S2, Supplementary Tables S1 – S6, and Supplementary Figures S1 – S2, which are referred to in the main text.

**Supplementary_File_1_Observatory_SamplingEvent_Image_Omics_DataMeta_data_re lease_001.xlsx**

Is referred to in the main text and contains metadata on observatories and sampling events for data_release_001 of ARMS-MBON, as well as links to image data and ENA accession numbers of genetic data for sampling events presented in this manuscript. It also contains info for Omics data on how sequence reads were demultiplexed.

**Supplementary_File_2_abundance_phyla_species_identified_data_release_001.xlsx**

Is referred to in the main text. The first three sheets contain abundances based on relative read numbers of identified phyla for the filtered and curated exploratory data set of all three marker genes. The last three sheets contain the number of species identified for each phylum (and class in case of ITS) for the three marker gene data sets. For the COI data set, taxonomic assignments below a confidence level of 0.8 were set as NA.

**Supplementary_File_3_Observatories_AMBI_RedList_WRiMS_species.xlsx**

Is referred to in the main text. Each sheet represents one of the 15 observatories and contains the identified species (with occurrences of at least two sequence reads) at each observatory which are registered as: i) very sensitive to disturbance in AMBI; ii) alien at the location of occurrence in WRiMS; and iii) Near Threatened, Vulnerable, Endangered or Critically Endangered in the Red Lists of the IUCN and HELCOM. Data of COI and 18S marker genes were pooled for each observatory.

**Supplementary_File_4_erroneus_sequences_removed.xlsx**

Is referred to in Supplementary Text S2. It contains two sheets, one each for the COI and 18S. Each sheets contains the ASVs/OTUs which were removed as potential contaminants based on their taxonomic classification, and the samples they occurred in with the respective read number.

