## Supplementary Material for "A long-term ecological research data set from the marine genetic monitoring programme ARMS-MBON 2018-2020"

### Supplementary Texts S1 to S2

#### Text S1: Bioinformatics processing of raw sequence data

Sequence data were processed with the Pipeline for Environmental DNA Metabarcoding Analysis, PEMA v.2.1.4 (Zafeiropoulos et al., 2020). PEMA currently supports the analysis of five marker genes (12S, 16S, 18S rRNA, COI, and ITS), and may also be used for other genes of interest if a taxonomic reference database is provided. PEMA consists of four main steps: (i) sequence pre-processing, (ii) OTU clustering or ASV inference, (iii) taxonomic assignment, and (iv) optionally the performance of biodiversity analysis based on the taxonomic inventory retrieved. It has been shown that parameter settings can lead to rather different outcomes (Brandt et al., 2021; Zafeiropoulos et al., 2020). Therefore, for comparison reasons, a fixed set of parameters was used for each marker gene and sequence data were processed separately for each sequencing run. All parameter files with their specific settings used as input for PEMA runs are available on the ARMS-MBON GitHub repository (see Supplementary Table S1 for respective link). For the 18S marker gene, sequences were clustered into operational taxonomic units (OTUs) using the VSEARCH v2.9.1 algorithm (Rognes et al., 2016) with a threshold of 0.97, while for ITS and COI, clustering was performed with Swarm v2 (Mahe et al., 2015), applying a threshold of d = 10 to infer amplicon sequence variants (ASVs). Note that PEMA initially defined the result of Swarm processing as inferred ASVs, i.e., sequences which differ by one or more nucleotides, which is now corrected to swarm-clusters (Hakimzadeh et al., 2023). We here use PEMA’s terminology for consistency reasons. Taxonomy was assigned to 18S OTUs and ITS ASVs with the CREST LCAClassifier v3.0 (Lanzè et al., 2012), using the PR2 v.4.13.0 (Guillou et al., 2013) and Unite v7.2 (Nilsson et al., 2018) databases, respectively.

Default settings were: i) minimum bit-score = 155; ii) LCA bitscore range = 2%; and iii) similarity cut-offs of 99%, 97%, 95%, 90%, 85% and 80% for the species, genus, family, order, class and phylum ranks, respectively. For COI sequences, taxonomic annotation was performed using the RDP classifier (Wang et al., 2007) with the MIDORI database v2.0 (Machida et al., 2017), and confidence values for each rank assignment were recorded, but no threshold was applied. Singletons (sequences with a total read abundance of one), OTUs/ASVs unclassified at domain level, and potential contaminant sequences (OTUs/ASVs that were more abundant in the negative control samples compared to actual samples) were removed. For OTUs/ASVs that were present in negative control samples in lower abundances than in actual samples, their corresponding read number in the negative controls was subtracted from their read number in actual samples. All bioinformatics analyses were supported by the High Performance Computing system of the Institute of Marine Biology, Biotechnology and Aquaculture of HCMR (Crete, Greece) (Zafeiropoulos et al., 2021).

**Text S2: Exploration of sequencing data**

The data from individual PEMA runs we provide on GitHub were merged for each marker gene and further curated to obtain a data set for visualisations and ecological assessments. As no confidence threshold was applied within PEMA for taxonomic assignments of COI ASVs (note that this is therefore also the case for the EurOBIS submission and users are urged to apply their own self-chosen cut-off), we excluded all rank assignments with a confidence value of below 0.8 for this marker gene. We further removed the sediment and plankton samples to solely assess the ARMS mobile and sessile data. Subsequently, we removed replicates of certain samples to reduce diversity inflation of the data set: i) where samples were re-sequenced and both versions of the respective sample pair remained in the data set, rarefaction curves, ASV/OTU count tables and taxonomic profiles of those samples were assessed and the version with higher diversity and/or better taxonomic resolution was kept; ii) for cases where samples remaining in the data set were preserved as duplicates in both EtOH and DMSO, only the DMSO sample was kept, as this is now the standard preservative used within ARMS-MBON; and iii) for the biological sample from the Roscoff (France) observatory which was processed as a technical duplicate (i.e., two replicates with the same preservative), rarefaction curves were generated and only the sample with the higher sequence richness and read count was kept.

Sequences with one of the following classifications were discarded as potential contaminants: *Homo sapiens*, *Canis lupus*, *Bos* spp. and *Zea mays*. Sequences classified as Insecta were removed if their lowest rank assignment was not listed in the World Register of Marine Species (WoRMS; Ahyong et al., 2024) as a marine and/or brackish taxon. See Supplementary File 4 for taxonomy and read abundances of removed sequences. One ASV in the ITS data identified as *Petrophila incerta* was incorrectly classified as the insect genus of the same name in the Unite database. We corrected its taxonomy to the *Petrophila* genus of the fungal Extremaceae family (Ascomycota), as this is a fungal species.

We determined the number and relative read abundance of unique phyla recovered through the application of the three marker genes, as well as the number of i) ASVs/OTUs classified with a Linnaean species name, ii) unique species identified, iii) species identified within each phylum, and iv) species shared between the data sets of the three marker genes. As the deposited taxonomy of different taxa does not necessarily follow the same Linnaean classification ranks within and across reference databases, some of the assigned taxonomies did not have a correct phylum level classification. For these cases, we manually determined the actual phylum classification of each infra-/subphylum, class, order, etc. present in our data set through a web-based search. We retrieved the correct phylum name from WoRMS; where this information could not be found in WoRMS, we relied on further scientific literature. In terms of alpha diversity, we assessed the observed ASV/OTU richness and the number of identified species (at the classification confidence threshold applied here) across observatories, as well as frequency distributions of these two parameters (i.e., re-occurrence of ASVs/OTUs and species identified across observatories).

We also assessed the influence of sampling effort on diversity variables. Here, we computed Spearman's correlation of sequencing depth (i.e., read number) and ARMS deployment duration (measured in days) versus ASV/OTU richness and the number of species identified in each sample. Furthermore, we computed Spearman's correlation of the number of ARMS units deployed and the number of samples included in the analysis post-curation versus ASV/OTU richness and the number of species identified at each of the 15 observatories. Where the correlation was statistically significant (i.e., $p < 0.05$) and moderate to strong (i.e., Spearman's $\rho > 0.4$), we performed analysis of simple linear regression to model the relationship between sampling effort predictor variables (i.e., sequencing depth measured as read numbers for sample-wise data and the number of ARMS units deployed and number of samples analysed for observatory-wise data) and dependent variables (i.e., ASV/OTU richness and number of species identified).

In order to test the application potential of the derived species observation data, we performed a scan against reference checklists for ecological key species. For this, we pooled COI and 18S data of species occurrences with at least two sequence reads and derived a list of species identified from each ARMS unit for each observatory. This list was then scanned against the following databases: i) AZTI's Marine Biotic Index (AMBI; Borja et al., 2000, 2019) for species very sensitive to disturbance; ii) the World Register of Introduced Marine Species (WRiMS; Costello et al., 2021, 2024) for species with alien status at the place of observation; and iii) the Red Lists of the International Union for Conservation of Nature (IUCN) and Baltic Marine Environment Protection Commission (Helsinki Commission, HELCOM) for species registered as Near Threatened, Vulnerable, Endangered or Critically Endangered. To this end, we used the web services provided by WoRMS. The AMBI and IUCN/HELCOM information were obtained using the WoRMS REST services (https://www.marinespecies.org/rest/; more specifically the call AphiaAttributesByAphiaID), while the WRiMS checks can be replicated using the Jupyter notebook on https://www.github.com/vliz-be-opsci/lw-iji-invasive-checker. We confirmed occurrences of red-listed species by scanning against known distribution in WoRMS and removed the *Pinna nobilis* occurrence from the Plymouth (UK) observatory, as this species is only known from the Mediterranean Sea and showed a low prevalence of only five reads in the respective sample.

Samples were tested for differences in alpha diversity among locations with varying degrees of anthropogenic influence (i.e., industrial, semi-industrial, low human influence (LHI), and protected; see Supplementary File S1 for influence category of each ARMS unit). After evaluating the deployment locations by consulting each network member, we identified these four categories appropriately describing the anthropogenic influence at each deployment site. Network members then classified each site according to the best-fitting category. For statistical comparison, samples with less than 5,000 reads were removed and the remaining samples rarefied to an equal sequencing depth of 5,000 reads without replacement to reduce diversity bias due to differences in sequencing depth. Given the relatively low number of remaining samples classified as "industrial" (i.e., n = 4 for COI, and n = 6 for 18S), these samples were grouped into one category ("industrial/semi-industrial") with samples classified as "semi-industrial". Mean and standard deviation (SD) of the two alpha diversity measures were calculated for samples of each influence type (i.e., three levels: industrial/semi-industrial, low human influence, and protected) and values were rounded to the nearest whole number. Data was subsequently checked for normality using the Shapiro-Wilk test. If data were normally distributed or log(1+x)-transformation resulted in normality (i.e., for $p > 0.05$), unidirectional analysis of variance (ANOVA) was applied to test for statistically significant differences between habitats. In case of significant differences (i.e., for $p < 0.05$), post-hoc Tukey's test was performed for pairwise comparisons. Where data was not normally distributed and transformation did not achieve normality, non-parametric Kruskal-Wallis rank sum test was applied (no statistical differences were revealed with this test, hence, no post-hoc test was performed).

As described above, all code used for exploratory analysis can be found at the dedicated GitHub repository (see Supplementary Table S1 for link). Analyses and data visualisation were performed in R v4.1.0 (R Core Team, 2021) via RStudio v2022.07.1 (RStudio Team, 2022) using packages of the tidyverse v1.3.1 collection (Wickham et al., 2019) and the packages Biostrings v2.60.2 (Pagès et al., 2020), phyloseq v1.36.0 (McMurdie & Holmes, 2013), vegan v2.6.2 (Oksanen et al., 2023), ggpubr v0.4.0 (Kassambara, 2020), grafify v4.0 (Shenoy, 2021), plyr v1.8.7 (Wickham, 2011), scales v1.3.0 (Wickham et al., 2023), egg v0.4.5 (Auguie, 2019), UpSetR v1.4.0 (Conway et al., 2017), xlsx v0.6.5 (Dragulescu & Arendt, 2020), writexl v1.5.0 (Ooms, 2024), and openxlsx v4.2.5 (Schauberger & Walker, 2021).

### Supplementary Tables S1 to S6

**Table S1.** Overview of ARMS-MBON project web pages, GitHub repositories and IMIS metadata records for taxonomic occurrences of data release 001 submitted to EurOBIS.

| **a) ARMS-MBON main webpages and GitHub documentation and data repositories** |
|---|
| *ARMS-MBON data landing page*<br>https://data.arms-mbon.org |
| *ARMS-MBON main GitHub page*<br>https://github.com/arms-mbon |
| *Documentation repository*<br>https://github.com/arms-mbon/documentation |
| *ARMS-MBON Handbook version applied for first sampling campaign*<br>https://github.com/arms-mbon/documentation/tree/main/armsmbon_handbook/old |
| *Molecular Standard Operating Procedures (MSOP)*<br>https://github.com/arms-mbon/documentation/tree/main/standard_operating_procedures |
| *All ARMS-MBON harvested metadata and analysis data organised in folders*<br>https://github.com/arms-mbon/data_workspace |
| *All ARMS-MBON quality-controlled observatory, sampling event, image and genetic metadata*<br>https://github.com/arms-mbon/data_workspace/tree/main/qualitycontrolled_data/combined |
| **b) *processing_batch1* repository** |

| |
|---|
| *Main page for all results files of PEMA bioinformatics processing of genetic data* <br> https://github.com/arms-mbon/data_workspace/tree/main/analysis_data/from_pema/processing_batch1 |
| *Parameter files used for PEMA runs* <br> https://github.com/arms-mbon/data_workspace/tree/main/analysis_data/from_pema/processing_batch1/parameter_files |
| *Fasta files with ASVs/OTUs resulting from PEMA processing* <br> https://github.com/arms-mbon/data_workspace/tree/main/analysis_data/from_pema/processing_batch1/fasta |
| *ASV/OTU tables and taxonomic assignments resulting from PEMA processing* <br> https://github.com/arms-mbon/data_workspace/tree/main/analysis_data/from_pema/processing_batch1/taxonomic_assignments |
| **c) *data_release_001* repository** |
| *data_release_001 main page* <br> https://github.com/arms-mbon/data_release_001/tree/main |
| *Info on observatories for which data was analysed for this data release* <br> https://github.com/arms-mbon/data_release_001/blob/main/ObservatoryData_release001.csv |
| *Info on sampling events and material samples* <br> https://github.com/arms-mbon/data_release_001/blob/main/SamplingeventData_release001.csv |
| *Download links for ARMS image data* <br> https://github.com/arms-mbon/data_release_001/blob/main/ImageData_release001.csv |
| *Info on amplicon sequencing data and corresponding ENA accession numbers* <br> https://github.com/arms-mbon/data_release_001/blob/main/OmicsData_release001.csv |
| **d) *analysis_release_001* repository** |
| *analysis_release_001 main page* <br> https://github.com/arms-mbon/analysis_release_001/tree/main |
| *Parameter files used for PEMA runs* <br> https://github.com/arms-mbon/analysis_release_001/tree/main/parameter_files |
| *Fasta files with ASVs/OTUs resulting from PEMA processing* <br> https://github.com/arms-mbon/analysis_release_001/tree/main/fasta |

| |
|---|
| *ASV/OTU tables and taxonomic assignments resulting from PEMA processing*<br>https://github.com/arms-mbon/analysis_release_001/tree/main/taxonomic_assignments |
| **d) *code_release_001* repository** |
| *Code used for exploratory data analysis*<br>https://github.com/arms-mbon/code_release_001 |
| **e) *IMIS metadata records for taxonomic occurrences submitted to EurOBIS*** |
| *COI*<br>https://www.vliz.be/en/imis?module=dataset&dasid=8357<br>*18S*<br>https://www.vliz.be/en/imis?module=dataset&dasid=8617<br>*ITS*<br>https://www.vliz.be/en/imis?module=dataset&dasid=8612 |

**Table S2.** Sampling effort and diversity measures per observatory for the COI and 18S marker genes. The number of samples analysed represents the number of samples remaining in the data set for each observatory after curation and filtering for data analysis. The number of reads equals the cumulative number of sequence reads in all samples used for data analysis. Number of species identified for COI was subject to the classification confidence threshold of 0.8 applied here.

| Observatory | No. of ARMS units deployed | COI | | | | 18S | | | |
|---|---|---|---|---|---|---|---|---|---|
| | | No. of samples analysed | No. of reads | ASV richness | No. of species identified | No. of samples analysed | No. of reads | OTU richness | No. of species identified |
| GulfOfPiran | 1 | 3 | 62946 | 3681 | 67 | 3 | 63188 | 567 | 13 |
| BelgiumCoast | 2 | 5 | 14811 | 2318 | 75 | 5 | 142944 | 265 | 18 |
| Bodo | 2 | 6 | 107908 | 2275 | 168 | 6 | 177301 | 923 | 20 |
| Gdynia | 2 | 6 | 52059 | 2058 | 67 | 6 | 108049 | 992 | 25 |
| TZS | 2 | 6 | 84961 | 5472 | 70 | 5 | 143828 | 636 | 18 |
| Getxo | 3 | 9 | 7497 | 795 | 63 | 9 | 13443 | 253 | 2 |
| Laeso | 3 | 8 | 4610 | 365 | 84 | 9 | 234634 | 1021 | 25 |
| Eilat | 4 | 12 | 17421 | 2197 | 60 | 12 | 17967 | 753 | 13 |
| Svalbard | 4 | 11 | 13996 | 2633 | 94 | 11 | 410560 | 1309 | 35 |
| Vigo | 4 | 12 | 79706 | 2618 | 113 | 10 | 184248 | 620 | 18 |
| Crete | 5 | 13 | 87425 | 2011 | 79 | 15 | 325088 | 943 | 33 |
| Limfjord | 5 | 15 | 175119 | 3179 | 130 | 15 | 151523 | 934 | 19 |
| Plymouth | 5 | 14 | 107742 | 4317 | 160 | 15 | 124524 | 1216 | 29 |
| Koster | 6 | 18 | 264774 | 5297 | 246 | 17 | 425795 | 2136 | 53 |
| Roscoff | 8 | 24 | 142485 | 4752 | 215 | 24 | 352153 | 1756 | 37 |

**Table S3.** Results of Spearman's correlation analyses for the association between parameters of sampling effort (i.e., deployment duration in days and sequencing depth measured as read number for sample-wise computations; number of deployed ARMS units and number of samples included in the analysis for observatory-wise computations) and diversity measures (i.e., ASV/OTU richness and number of species identified).

| Relationship investigated for Pearson's correlation | COI | 18S |
|---|---|---|
| **Per sample** | | |
| ASV/OTU richness vs. sequencing depth (i.e., read number) | $S$ = 283590<br>$p$ < .001<br>Spearman's $\rho(160)$ = .60<br>n = 162 | $S$ = 114054<br>$p$ < .001<br>Spearman's $\rho(160)$ = .84<br>n = 162 |
| Number of identified species vs. sequencing depth (i.e., read number) | $S$ = 159033<br>$p$ < .001<br>Spearman's $\rho(160)$ = .78<br>n = 162 | $S$ = 132478<br>$p$ < .001<br>Spearman's $\rho(160)$ = .81<br>n = 162 |
| ASV/OTU richness vs. deployment duration in days | $S$ = 811528<br>$p$ = .065<br>Spearman's $\rho(160)$ = -.15<br>n = 162 | $S$ = 654335<br>$p$ = .333<br>Spearman's $\rho(160)$ = .08<br>n = 162 |
| Number of identified species vs. deployment duration in days | $S$ = 658469<br>$p$ = .371<br>Spearman's $\rho(160)$ = .07<br>n = 162 | $S$ = 613801<br>$p$ = .090<br>Spearman's $\rho(160)$ = .13<br>n = 162 |
| **Per observatory** | | |
| ASV/OTU richness vs. number of ARMS units deployed | $S$ = 414.51<br>$p$ = .350<br>Spearman's $\rho(13)$ = .26<br>n = 15 | $S$ = 193.73<br>$p$ < .008<br>Spearman's $\rho(13)$ = .65<br>n = 15 |
| Number of identified species vs. number of ARMS units deployed | $S$ = 224.46<br>$p$ = .018<br>Spearman's $\rho(13)$ = .60<br>n = 15 | $S$ = 209.14<br>$p$ = .012<br>Spearman's $\rho(13)$ = .63<br>n = 15 |
| ASV/OTU richness vs. number of samples remaining in the data set for ecological analysis | $S$ = 402.29<br>$p$ = .309<br>Spearman's $\rho(13)$ = .28<br>n = 15 | $S$ = 173.58<br>$p$ = .004<br>Spearman's $\rho(13)$ = .69<br>n = 15 |
| Number of identified species vs. number of samples remaining in the data set for ecological analysis | $S$ = 230.23<br>$p$ = .021<br>Spearman's $\rho(13)$ = .59<br>n = 15 | $S$ = 202.85<br>$p$ = .011<br>Spearman's $\rho(13)$ = .64<br>n = 15 |

**Table S4.** Results of linear regression analyses for the relationship between parameters of sampling effort (i.e., deployment duration in days and sequencing depth measured as read number for sample-wise computations; number of deployed ARMS units and number of samples included in the analysis for observatory-wise computations) and diversity measures (i.e., ASV/OTU richness and number of species identified). Note that regression analysis was only performed for associations where Pearson's correlation was significant (i.e., $p < .05$) and moderate to strong (i.e., Pearson's $r > .4$), see Table S2. Significance codes: 0 - ***; 0.001 - **; 0.01 - *.

<table>
<tr><th colspan="2">Per sample</th></tr>
<tr><th>Marker gene</th><th>ASV/OTU richness vs. sequencing depth (i.e., read number)</th></tr>
<tr><td>COI</td><td>Residuals:<br>Min 1Q Median 3Q Max<br>-651.38 -188.66 -75.57 71.04 1566.87<br><br>Coefficients:<br>Estimate Std. Error t value Pr(>|t|)<br>(Intercept) 2.209e+02 2.809e+01 7.865 5.16e-13 ***<br>reads 1.377e-02 1 .997e-03 6.898 1.16e-10 ***<br>---<br>Residual standard error: 301.6 on 160 degrees of freedom<br>Multiple R-squared: 0.2292, Adjusted R-squared: 0.2244<br>F-statistic: 47.58 on 1 and 160 DF, p-value: 1.163e-10</td></tr>
<tr><td>18S</td><td>Residuals:<br>Min 1Q Median 3Q Max<br>-368.34 -76.32 -19.77 43.07 644.21<br><br>Coefficients:<br>Estimate Std. Error t value Pr(>|t|)<br>(Intercept) 1.019e+02 1.082e+01 9.418 < 2e-16 ***<br>reads 2.261e-03 2.817e-04 8.026 2.04e-13 ***<br>---<br>Residual standard error: 122.2 on 160 degrees of freedom<br>Multiple R-squared: 0.287, Adjusted R-squared: 0.2826<br>F-statistic: 64.41 on 1 and 160 DF, p-value: 2.042e-13</td></tr>
<tr><th>Marker gene</th><th>Number of identified species vs. sequencing depth (i.e., read number)</th></tr>
<tr><td>COI</td><td>Residuals:<br>Min 1Q Median 3Q Max<br>-40.077 -12.729 -2.018 8.236 51.231<br><br>Coefficients:<br>Estimate Std. Error t value Pr(>|t|)<br>(Intercept) 1.858e+01 1.513e+00 12.282 < 2e-16 ***<br>reads 7.361e-04 1.075e-04 6.844 1.55e-10 ***<br>---<br>Residual standard error: 16.25 on 160 degrees of freedom<br>Multiple R-squared: 0.2265, Adjusted R-squared: 0.2216<br>F-statistic: 46.85 on 1 and 160 DF, p-value: 1.552e-10</td></tr>
<tr><td>18S</td><td>Residuals:</td></tr>
</table>

| | |
|---|---|
| | *Min 1Q Median 3Q Max*<br>*-16.1862 -2.3383 -0.8963 1.2926 27.0470*<br><br>*Coefficients:*<br>*Estimate Std. Error t value Pr(>\|t\|)*<br>*(Intercept) 2.666e+00 3.768e-01 7.076 4.4e-11 ****<br>*reads 1.044e-04 9.805e-06 10.646 < 2e-16 ****<br>*---*<br>*Residual standard error: 4.253 on 160 degrees of freedom*<br>*Multiple R-squared: 0.4146, Adjusted R-squared: 0.411*<br>*F-statistic: 113.3 on 1 and 160 DF, p-value: < 2.2e-16* |
| **Per observatory** | |
| **Marker gene** | **ASV/OTU richness vs. number of ARMS units deployed** |
| 18S | *Residuals:*<br>*Min 1Q Median 3Q Max*<br>*-558.04 -264.98 12.52 255.85 736.30*<br><br>*Coefficients:*<br>*Estimate Std. Error t value Pr(>\|t\|)*<br>*(Intercept) 222.38 216.11 1.029 0.32225*<br>*arms 196.22 52.11 3.766 0.00236 ***<br>*---*<br>*Residual standard error: 364.5 on 13 degrees of freedom*<br>*Multiple R-squared: 0.5217, Adjusted R-squared: 0.4849*<br>*F-statistic: 14.18 on 1 and 13 DF, p-value: 0.002357* |
| **Marker gene** | **ASV/OTU richness vs. number of samples analysed** |
| 18S | *Residuals:*<br>*Min 1Q Median 3Q Max*<br>*-585.72 -283.19 -10.09 230.12 780.79*<br><br>*Coefficients:*<br>*Estimate Std. Error t value Pr(>\|t\|)*<br>*(Intercept) 257.68 209.29 1.231 0.2400*<br>*samples 64.56 17.29 3.734 0.0025 ***<br>*---*<br>*Residual standard error: 366.1 on 13 degrees of freedom*<br>*Multiple R-squared: 0.5175, Adjusted R-squared: 0.4804*<br>*F-statistic: 13.94 on 1 and 13 DF, p-value: 0.002502* |
| **Marker gene** | **Number of identified species vs. number of ARMS units deployed** |
| COI | *Residuals:*<br>*Min 1Q Median 3Q Max*<br>*-61.274 -18.660 -5.531 11.597 92.954*<br><br>*Coefficients:*<br>*Estimate Std. Error t value Pr(>\|t\|)*<br>*(Intercept) 31.561 26.348 1.198 0.25235* |

| | |
|---|---|
| | *arms 21.743 6.353 3.422 0.00454 ***<br>---<br>*Residual standard error: 44.44 on 13 degrees of freedom*<br>*Multiple R-squared: 0.474, Adjusted R-squared: 0.4335*<br>*F-statistic: 11.71 on 1 and 13 DF, p-value: 0.004542* |
| 18S | *Residuals:*<br>*Min 1Q Median 3Q Max*<br>*-18.832 -5.745 1.305 4.030 19.755*<br><br>*Coefficients:*<br>*Estimate Std. Error t value Pr(>\|t\|)*<br>*(Intercept) 8.420 5.874 1.433 0.1753*<br>*arms 4.138 1.416 2.921 0.0119 **<br>---<br>*Residual standard error: 9.907 on 13 degrees of freedom*<br>*Multiple R-squared: 0.3963, Adjusted R-squared: 0.3499*<br>*F-statistic: 8.535 on 1 and 13 DF, p-value: 0.01191* |
| **Marker gene** | **Number of identified species vs. number of samples analysed** |
| COI | *Residuals:*<br>*Min 1Q Median 3Q Max*<br>*-61.800 -17.355 -7.578 9.644 91.533*<br><br>*Coefficients:*<br>*Estimate Std. Error t value Pr(>\|t\|)*<br>*(Intercept) 31.134 24.775 1.257 0.23098*<br>*samples 7.555 2.052 3.681 0.00277 ***<br>---<br>*Residual standard error: 42.87 on 13 degrees of freedom*<br>*Multiple R-squared: 0.5104, Adjusted R-squared: 0.4728*<br>*F-statistic: 13.55 on 1 and 13 DF, p-value: 0.002766* |
| 18S | *Residuals:*<br>*Min 1Q Median 3Q Max*<br>*-19.496 -4.533 1.773 3.553 20.967*<br><br>*Coefficients:*<br>*Estimate Std. Error t value Pr(>\|t\|)*<br>*(Intercept) 9.6417 5.7948 1.664 0.1200*<br>*samples 1.3171 0.4787 2.751 0.0165 **<br>---<br>*Residual standard error: 10.14 on 13 degrees of freedom*<br>*Multiple R-squared: 0.368, Adjusted R-squared: 0.3194*<br>*F-statistic: 7.57 on 1 and 13 DF, p-value: 0.01649* |

**Table S5.** Number of identified species (with occurrences of at least two sequence reads) across observatories listed in four different databases. Data of COI and 18S marker genes were pooled for each observatory. AMBI – species registered as very sensitive to disturbance in AZTI's Marine Biotic Index; Borja *et al*., 2000, 2019); WRiMS – species registered as alien at the location of occurrence in the World Register of Introduced Marine Species (Costello *et al*., 2014); IUCN/HELCOM Red List – species registered as Near Threatened, Vulnerable, Endangered or Critically Endangered in the Red Lists of the International Union for the Conservation of Nature and the Baltic Marine Environment Protection Commission (Helsinki Commission). Observatories are ordered from top to bottom by increasing number of ARMS units deployed (n = 1 for GulfofPiran, n = 8 for Roscoff).

| Observatory | AMBI | WRiMS | IUCN/HELCOM Red List |
|---|---|---|---|
| GulfOfPiran | 15 | 3 | 0 |
| BelgiumCoast | 17 | 4 | 0 |
| Bodo | 17 | 5 | 0 |
| Gdynia | 2 | 6 | 0 |
| TZS | 4 | 5 | 0 |
| Getxo | 8 | 0 | 0 |
| Laeso | 13 | 0 | 0 |
| Eilat | 7 | 0 | 0 |
| Svalbard | 11 | 0 | 0 |
| Vigo | 16 | 1 | 0 |
| Crete | 7 | 4 | 0 |
| Limfjord | 13 | 8 | 0 |
| Plymouth | 20 | 3 | 1 |
| Koster | 37 | 6 | 2 |
| Roscoff | 23 | 4 | 2 |

**Table S6.** Mean, standard deviation and results of statistical tests for differences in ASV/OTU richness and the number of species identified among habitats of varying degrees of anthropogenic influence for the COI and 18S data. Samples with less than 5,000 sequence reads were removed prior to the analysis and all remaining samples were rarefied to an even sequencing depth of 5,000 reads.

| **Statistical measure/test** | **COI** | |
|---|---|---|
| | **ASV richness** | **No. of species identified** |
| Mean ± standard deviation | Mean, SD<br>Industrial/Semi-industrial: 259.79, 200.10<br>Low Human Influence: 216.35, 89.98<br>Protected: 260.00, 172.93 | Mean, SD<br>Industrial/Semi-industrial: 29.00, 12.80<br>Low Human Influence: 26.24, 7.55<br>Protected: 38.09, 14.96 |
| Shapiro-Wilk test | W = 0.89085, p < 0.001<br>log(1+x)-transformed data: W = 0.98638, p = 0.7142 | W = 0.97576, p = 0.2479 |
| ANOVA | log(1+x)-transformed data:<br>ANOVA, Df, Sum Sq, Mean Sq, F value, Pr(>F)<br>Habitat, 2, 0.234, 0.1171, 0.241, 0.787<br>Residuals, 60, 29.2, 0.4867 | ANOVA, Df, Sum Sq, Mean Sq, F value, Pr(>F)<br>Habitat, 2, 1577, 788.3, 5.043, 0.00945<br>Residuals, 60, 9379, 156.3 |
| **Statistical measure/test** | **18S** | |
| | **OTU richness** | **No. of species identified** |
| Mean ± standard deviation | Mean, SD<br>Industrial/Semi-industrial: 123.00, 77.90<br>Low Human Influence: 136.05, 52.94<br>Protected: 165.22, 83.72 | Mean, SD<br>Industrial/Semi-industrial: 5.35, 4.50<br>Low Human Influence: 5.81, 3.89<br>Protected: 6.65, 3.01 |
| Shapiro-Wilk test | W = 0.96492, p = 0.06572 | W = 0.95471, p = 0.01963<br>log(1+x)-transformed data: W = 0.93212, p = 0.001658 |
| ANOVA/ Kruskal-Wallis | ANOVA, Df, Sum Sq, Mean Sq, F value, Pr(>F)<br>Habitat, 2, 20349, 10174, 1.906, 0.157<br>Residuals, 61, 325555, 5337 | Kruskal-Wallis chi-squared = 2.5433, df = 2, p = 0.2804 |

### Supplementary Figures S1 to S2

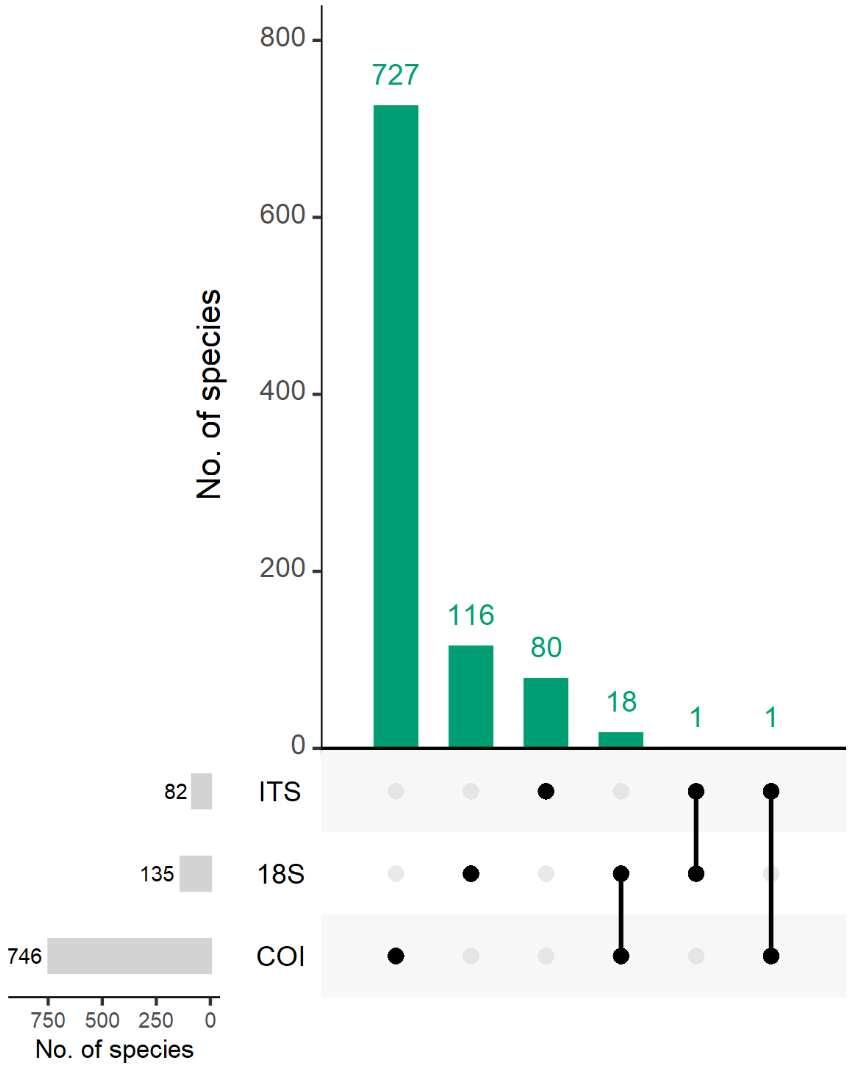


**Figure S1.** UpSet plot of the number of species identified using the three marker genes COI, 18S, and ITS. Green bars display the number of species identified (at the given confidence threshold applied here) that are shared across the three data sets. The matrix below the bar plot shows which combination of marker genes correspond to each bar. Bars on the left represent the total number of species identified in each marker gene's data set. No species were common to all three data sets.

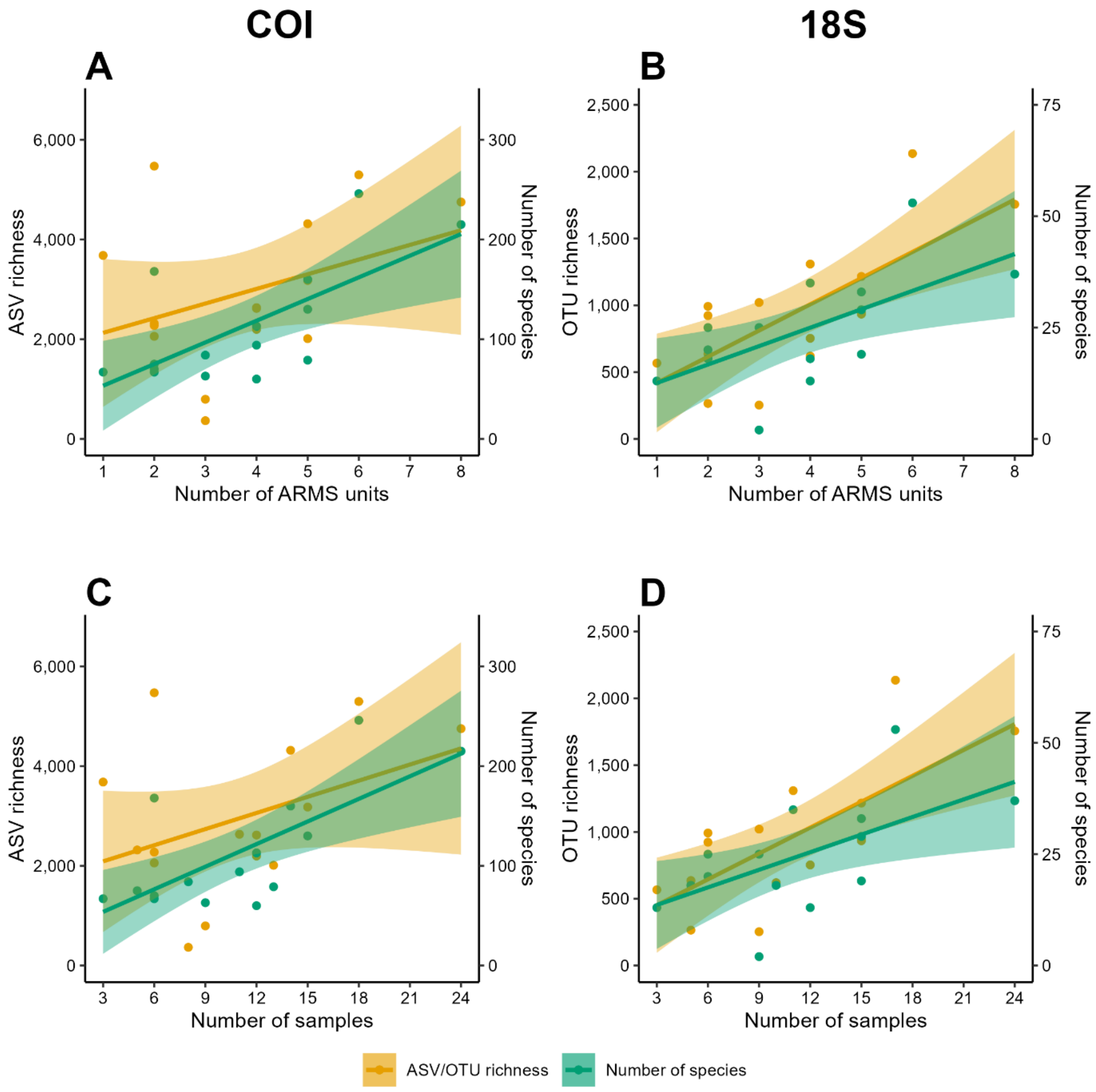


**Figure S2.** Relationship of ASV/OTU richness (yellow) and the number of species identified (green) for COI (**A**) and 18S (**B**) versus the number of ARMS units deployed at each observatory; and relationship of ASV/OTU richness (yellow) and the number of species identified (green) for COI (**C**) and 18S (**D**) versus the number of samples remaining in the data sets after curation for each observatory. Solid lines represent linear regression for ASV/OTU richness (yellow) and the number of species identified (green), shaded areas represent the corresponding 95% confidence intervals. Note that no significant linear correlation was found for COI ASV richness versus both sampling effort parameters in both **A** and **C**.
